# Habitat patch size and tree species richness shape the bird community in urban green spaces of rapidly urbanizing region of India

**DOI:** 10.1101/2020.10.23.348391

**Authors:** Monica Kaushik, Samakshi Tiwari, Kumari Manisha

## Abstract

Rapid urbanization and associated biodiversity loss is rampant globally but especially a cause of concern for developing countries. However, numerous studies investigating the role of urban green spaces have established their key role in conserving larger suites of species in urban area. Yet our knowledge is lopsided due to lag in research in developing countries. We examined how landscape and local scale features of urban green spaces influence bird species richness, density, fine-foraging guild richness and composition during breeding and non-breeding season. This is the first study of this nature in one the Himalayan states of India. We quantified landscape level variables in the 250m buffer around 18 urban green spaces. We sampled vegetation and bird community during breeding and non-breeding season through 52 intensive sampling point spread across 18 urban green spaces. Size of the urban green space at landscape level and tree richness at the local scale emerged as important predictor variables influencing bird species richness, density and richness of imperiled insectivorous guild across seasons. Urban green spaces within education institutions and offices experiencing much less management supported higher bird richness and density whereas city parks were the most species poor. Community composition was affected more strongly by built-up cover and barren area in the matrix and also by tree species richness within urban green spaces. City planners should focus on establishing larger city parks during design stage whereas biodiversity potential of the existing urban green spaces could be enhanced by selecting native tree and shrub species to increase overall habitat complexity.

## Introduction

Urban expansion is one of the biggest threats to biodiversity (Kang et al., 2015).In 2018, 55% of the worlds’ population was living in urban areas, which is expected to increase to 68% by 2050 (DESA, 2018). A sizeable amount of this expansion is expected from developing countries like India, China, and Nigeria. Urban areas are characterized by a mix of variety of grey and green spaces accommodating a large suite of common and highly plastic species. However, urban areas are also inhabited by few threatened species(Ives et al., 2016). Both common and threatened species play significant role in urban ecosystem functioning and provides multitude of ecosystem services. For example, in an experimental study conducted across three towns of UK reported higher amount of carcass removal in the presence of three urban vertebrate scavengers than in their absence (Inger et al., 2016).Varying in size and shape green spaces in urban areas ranges from city parks, remnant forest patches, golf courses to cemeteries act as hotspots of biodiversity (Gallo et al., 2017; Wurth et al., 2020). Variety of green habitats in urban areas covered partially or completely by any type of vegetation under private or public ownership are collectively known as urban green spaces.

In past decade, urban green spaces have received much required attention as a conservation tool for urban biodiversity. Urban green spaces can support endemic native species (Carbó-Ramírez & Zuria, 2011), mitigate urban heat island effect (Park et al., 2017; Xiao et al., 2018), ensure mental wellbeing of the visitors (Carrus et al., 2015) and prevent “extinction of experience” in human population residing in urban areas (Soga & Gaston, 2016).Studies focusing on habitat characteristics of urban greenspaces can improve biodiversity conservation potential(Aronson et al., 2017).

Previous studies have investigated the habitat features of the greenspaces largely at patch scale. Patch size emerges as an universal predictors across studies that improve biodiversity potential of greenspaces, conforming species-area relationship in urban ecosystem (Chamberlain et al., 2007; Dale, 2018; La Sorte et al., 2020; Matthies et al., 2015; Nielsen et al., 2014). Other than size of the park, habitat diversity within the urban green space and its age also positively influence the biodiversity (Zivanovic & Luck, 2016). Degree of connectivity among urban green spaces increases richness by allowing immigration of species from source habitats to other potential habitat (Braaker et al., 2017; Shanahan et al., 2011).

Urban green spaces are nested in varied matrix of habitat types that ranges from completely urban to a remnant forest patches. These habitats surrounding habitats, also known as matrix, can substantially influence the species richness and composition within the greenspaces. For example, higher proportion of “built-up” area in the matrix negatively affects the richness of bird species of the urban green spaces at community (Murgui, 2009) and guild level (Amaya-Espinel et al., 2019; Chamberlain et al., 2007; Fischer et al., 2016; Pellissier et al., 2012). Matrix with no or low management interventions such as fallow land or abandoned successional habitats often provide distinct resources and thereby elevate species richness of certain taxa (Melliger et al., 2017).

At the local scale, habitat heterogeneity within the urban green spaces in form of vegetation structure and complexity increases the richness and diversity of multiple taxa (Kang et al., 2015; Nielsen et al., 2014). Additionally, increase in tree and shrub diversity support faunal diversity at the local scale (Nielsen et al., 2014). Shrub cover could have different effects on richness depending on the focal taxa. Increasing shrub cover especially in highly urbanized matrix improved richness of highly imperiled insectivorous bird taxa (Pellissier et al., 2012) but reduced bee richness by reducing their nesting resources (Banaszak-Cibicka et al., 2016).Information on habitat features that improve the biodiversity potential of urban green spaces could be used by the urban planner and managers at design and maintenance stages of urban greening projects (Callaghan et al., 2018).

In this study we investigated how habitat features of urban green spaces at landscape and local scale affects the bird community and fine-foraging guilds during breeding and non-breeding seasons. Additionally, we investigated if bird species composition varies across urban green spaces and if so, which factors are responsible for the differences. We selected birds owing to the ease of quantification as well as their property of being a good surrogate of overall biodiversity (Eglington et al., 2012). Birds are also important ecosystem service providers especially in tropical countries where majority of plants depend on bird-mediated seed dispersal (Sekercioglu et al., 2016; Whelan et al., 2008), preventing crop damage by arthropods control (Maas et al., 2016) and pollination (S. H. Anderson et al., 2016). Therefore, conservation of birds through urban green spaces ensures maintenance of diverse ecosystem services provided by them in urban areas. Our aim was to examine whether and how urban green spaces can be planned and managed to improve species richness, density, and guild richness in urban ecosystem.

## Materials and methods

### Study Area

We carried out this study in Dehradun city (30.3165° N, 78.0322° E) which is the capital of the northern state, Uttarakhand, India. It is located at the foothills of Himalaya flanked by two important rivers, Yamuna and Ganga. Dehradun is a valley spread across an area of 3088 km^2^ with moderate variation in elevation (410m-700m). The city is characterized by mild weather throughout the year, but winter’s temperature could be as low as 0-1°C and the maximum temperature in summers could be as high as 40°C. However, maximum temperature during summer is increasing. For example, in 2019 a maximum temperature of 44°C was recorded for the first time in the month of May. The area receives an average annual rainfall of 2073 mm, largely during the monsoon season (July-August).

Uttarakhand state was carved out from the Uttar Pradesh in year 2000 and Dehradun was designated its capital. Changed political status resulted in push towards infrastructural and developmental activities at the cost of the agricultural, forest and open areas. Between the years 2001 and 2011 Dehradun experienced rapid population growth (https://www.census2011.co.in/census/district/578-dehradun.html). Though Dehradun has 64 city parks (http://smartcities.gov.in/upload/uploadfiles/files/Annexures_Dehradun.pdf), most of these are small parks constructed within residential colonies. Majority of urban green spacesin Dehradun – and other cities within India –are in the form of personal gardens, fruit orchards, tea gardens, tree belts along *nallahs* and reserved forests. In recent years, green spaces in Dehradun have shrunk due to increasing built-up for residential, commercial, and industrial purposes(Dutta et al., 2015). However, abutting Himalayan foothills Dehradun harbors 42% (567 of 1338) of the avifaunal diversity of India and 82% (567 of 688) of Uttarakhand state (www.ebird.org/India). Different habitats within the city provide safe breeding and wintering ground to the summer and winter migratory birds(Mohan, 2007).

### Study site selection

We selected sites across a gradient of urban green space size using satellite imagery of Google Earth (*Google Earth Pro*, 2018). While selecting sites we made sure that the sites were spatially distributed evenly across the city. Sites were visited for ground-truthing to assess the suitability in terms of accessibility and vegetation type. We avoided orchard of cash crops which generally lack shrub layer and are not open to public. We did choose one old tea plantation due to its large size, presence of native trees and continuous reporting of rare birds (e.g., Himalayan Griffon *Gyps himalayensis*, Yellow-eyed Babbler *Chrysomma sinense*). Out of 28 urban green spaces identified using Google Earth imagery, 18 sites were shortlisted for the study (Figure 1). Using ArcGIS (version 10.6) we measured the area of selected sites. We quantified the matrix composition around each urban green spaces within a buffer of 250 m using ArcGIS (version 10.6) software. The following landuse types-agricultural field, green cover (including woodland), open (scrubland) areas, water cover, built-up and barren were digitized using polygon tool of Google Earth and later quantified for their extent using the ArcGIS (version 10.6).

**Figure 1:**
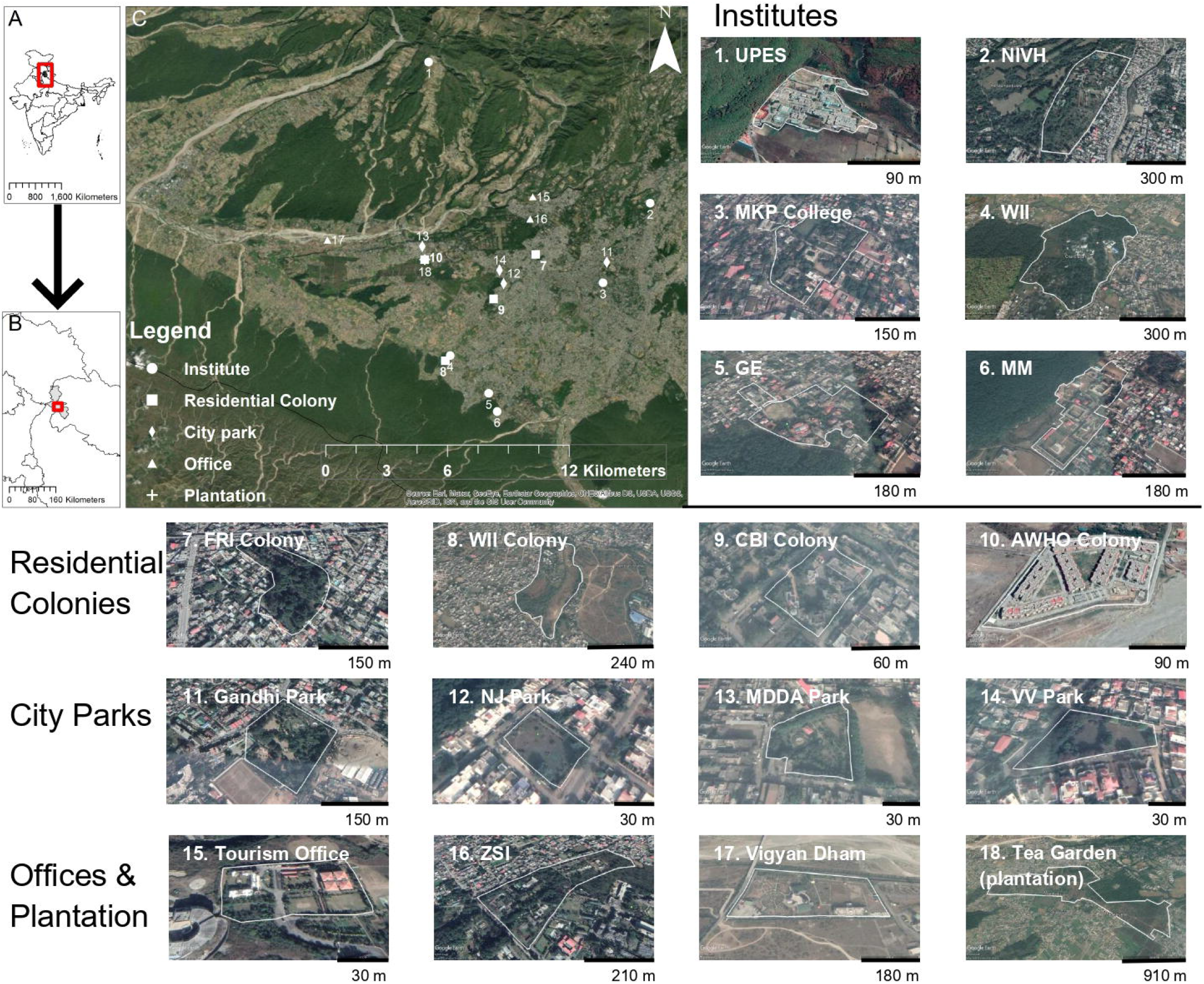
Map of study area showing 18 urban green spaces selected for bird and vegetation sampling in Dehradun, Uttarakhand, India. The number allotted to each urban green space represents its location on the map.

### Quantification of habitat structure and composition

Each urban green space was divided into sampling grids of 200m and the centroids of the grid were selected for intensive vegetation and bird sampling. At each plot we recorded structural and compositional features of the vegetation by quantifying the trees and shrubs within concentric plots of 20m and 5m radius, respectively. For structural features of the tree layer we recorded girth at breast height, total and bole height and canopy spread in two perpendicular axes. Bole and total height of the tree was quantified using an altimeter. For shrub structural features we recorded average height for each shrub species and its spread within 5m radius plot. We recorded each tree and shrub to species level with the help of available field guides (Kanjilal & Gupta, 1979)

### Sampling bird community

We sampled bird community using the variable radius point transect method centered on the vegetation sampling plots. We choose point transects for sampling birds as well-spaced point transects could provide finer information than line-transects about the bird-habitat relationship if habitat parameters are quantified around the points (Bibby et al., 2000).

All the point transects were conducted by only a single observer in one season (ST: non-breeding season and KM: breeding season) to avoid observer bias and all species seen or heard were recorded at the point. The observer also recorded the radial distance of each observation using a laser rangefinder. Bird sampling was carried out in morning hours (6:00 am – 9:00 am) during breeding (March-May) and non-breeding season (September-December). Each site was visited four times each within breeding and non-breeding season. Species were recorded for 7 minutes after 3 minutes of acclimatization time. To capture the maximum species variation within a season, each site was revisited after a week. The order of visiting the points was reversed on each morning to negate the bias due to flushing of birds by observer. A total of 416 (52 points x 4 times x 2 seasons) variable radius point transects were undertaken during the study period.

### Data analysis

For each urban green space, we estimated the richness for bird, tree and shrub species using package vegan (Oksanen et al., 2013) within R platform(R Core Team, 2019). We estimated bird species richness separately for each season using first-order jackknife richness estimator. Overall bird density for each urban green space was estimated using the program DISTANCE 7.3 (Thomas et al., 2010).

We used linear modeling approach to evaluate the relationship between landscape and local scale variables on bird species richness, overall bird density and richness of fine-foraging guild. We categorized birds into their fine-foraging guilds using the information provided by Mohan (2007) in the same site. We used generalized linear models with Poisson family for modeling the guild species richness. Considering the differences in spatial scales, we built models separately for landscape and local scale variables (Electronic supplementary material A, B, C and D).

Area of urban green space was log transformed for all analysis. We built models with only uncorrelated variables and selected the best model through an information criterion model selection approach (Burnham & Anderson, 2010). We used Akaike information criterion for small sample sizes (AICc) for model selection since the ratio of sample size (n) and number of parameters (K) was small (i.e., <40;(Burnham and Anderson 2010)). The model with the lowest AICc value and within 2 ΔAICc was selected as the best model(s). To estimate model coefficients, we used model averaging whenever there were more than one models within 2 ΔAICc values. Model averaging was performed using package MuMIn in R(Barton & Barton, 2015). We estimated the back transformed estimate and standard error of variables in the best model using package *arm* (Gelman et al., 2018).

We used Non-Metric Multidimensional Scaling (NMDS) to explore differences in bird species composition across each urban green space and the associated landscape and local-scale variables. We choose Bray-curtis dissimilarity index, which works well with the abundance data (M. Anderson, 2001). Rare and vagrant species seen only once during the study period were removed for performing this analysis. We explored the relationship between NMDS axis and the habitat covariates using the function *envfit* in package vegan. We used *adonis* test to explore if the bird species composition varied with the size and type of the urban green space. All statistical analyses were performed using program the R version 3.6.0 (R Core Team, 2019) and graphical visualization were created using ggplot2 (Wickham, 2016).

## Result

### Habitat characterization of the urban green spaces

We selected 18 urban green spaces of which six were educational institutions, four city parks, four residential complex, three offices parks and one old abandoned tea plantation. The area of urban green spaces varied from 0.3ha to 224 ha (Table 1), where abandoned tea plantation was the largest urban green spaces. The urban matrix around urban green spaces had relatively higher proportion of “built-up” than other land use types (Table 1). The second most abundant landuse type in the matrix was “green cover” that varied from 5.96 % to 60% (Table 1). Agricultural area was the least dominant landuse type in the matrix and ranged between 0 to 16% (Table 1). We recorded a total of 92 trees species and 112 shrubs species from the entire study area.

**Table 1:**
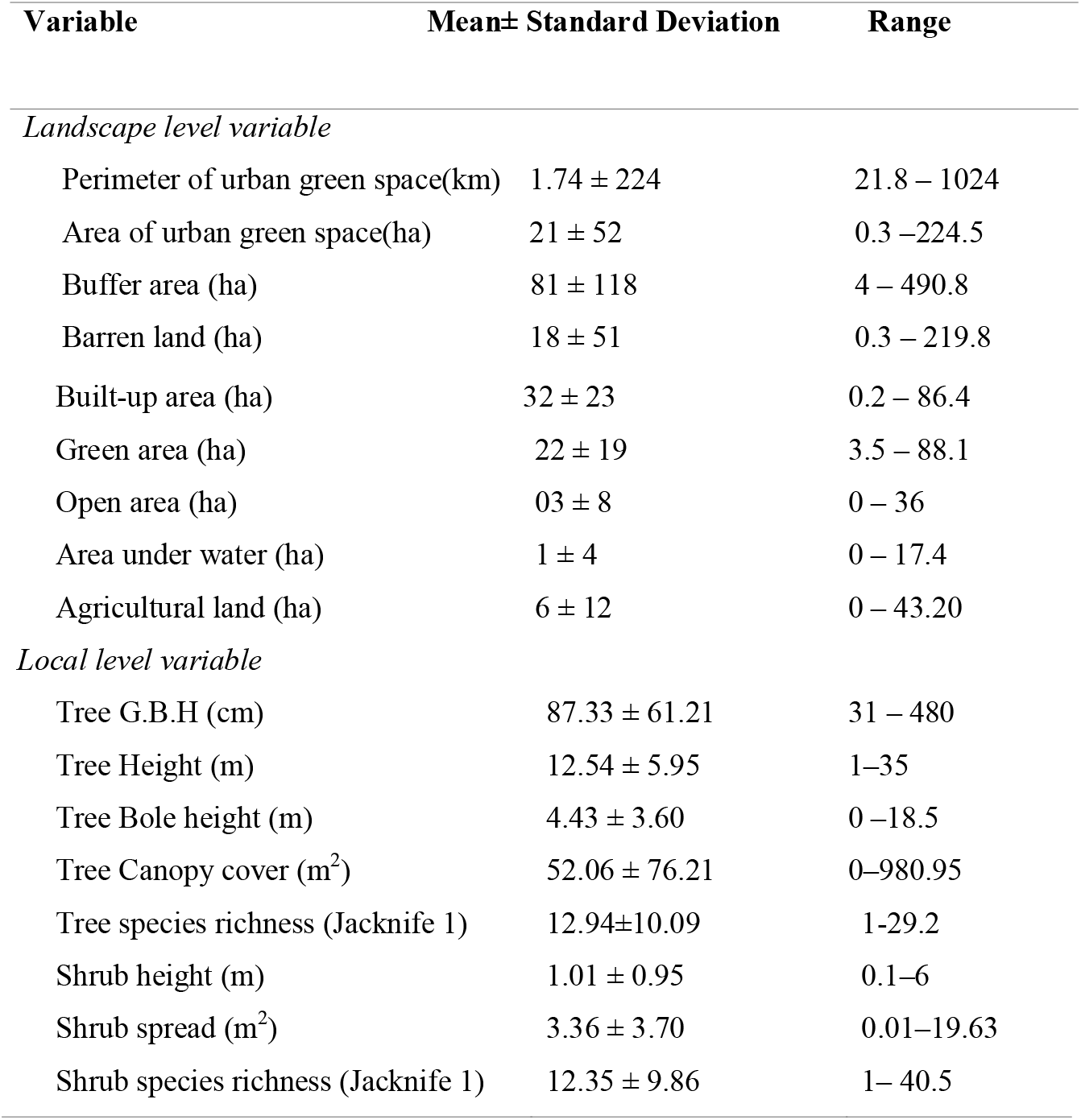
Average value of landscape and local scale variables across 18 urban green spaces of Dehradun, Uttarakhand, India.

| Variable | Mean± Standard Deviation | Range |
| --- | --- | --- |
| *Landscape level variable* |  |  |
| Perimeter of urban green space(km) | 1.74 ± 224 | 21.8 – 1024 |
| Area of urban green space(ha) | 21 ± 52 | 0.3 – 224.5 |
| Buffer area (ha) | 81 ± 118 | 4 – 490.8 |
| Barren land (ha) | 18 ± 51 | 0.3 – 219.8 |
| Built-up area (ha) | $32 \pm 23$ | 0.2 – 86.4 |
| Green area (ha) | $22 \pm 19$ | 3.5 – 88.1 |
| Open area (ha) | $03 \pm 8$ | 0 – 36 |
| Area under water (ha) | $1 \pm 4$ | 0 – 17.4 |
| Agricultural land (ha) | $6 \pm 12$ | 0 – 43.20 |
| *Local level variable* |  |  |
| Tree G.B.H (cm) | $87.33 \pm 61.21$ | 31 – 480 |
| Tree Height (m) | $12.54 \pm 5.95$ | 1–35 |
| Tree Bole height (m) | $4.43 \pm 3.60$ | 0 –18.5 |
| Tree Canopy cover (m<sup>2</sup>) | $52.06 \pm 76.21$ | 0–980.95 |
| Tree species richness (Jackknife 1) | $12.94 \pm 10.09$ | 1-29.2 |
| Shrub height (m) | $1.01 \pm 0.95$ | 0.1–6 |
| Shrub spread (m<sup>2</sup>) | $3.36 \pm 3.70$ | 0.01–19.63 |
| Shrub species richness (Jackknife 1) | $12.35 \pm 9.86$ | 1– 40.5 |

### Bird species richness and density

A total of 139 (4399 detections) species were recorded during the study period covering breeding and non-breeding season. Like other studies from this region conducted in natural forest(Kaushik, 2016)and urban forests (Mohan, 2007), bird species richness was higher during the breeding (123 species) than the non-breeding season (103 species) (Figure 2a). Older government institutes for education and research had the highest bird species richness whereas city parks had the lowest richness, consistently across breeding and non-breeding season. Overall bird density per hectare varied from 11.54_Mean_ ± 10.43%_cv_ to 143.02_Mean_ ± 19.36 %_cv_ during breeding season and 17.84_Mean_± 20.44%_cv_ to 154.83_Mean_ ± 16.99%_cv_ during non-breeding season. Urban green spaces within institutes and residential complexes had higher density during the breeding season than non-breeding season (Figure 2b). City park exhibited a high variation in bird density during the breeding season than non-breeding season.

**Figure 2:**
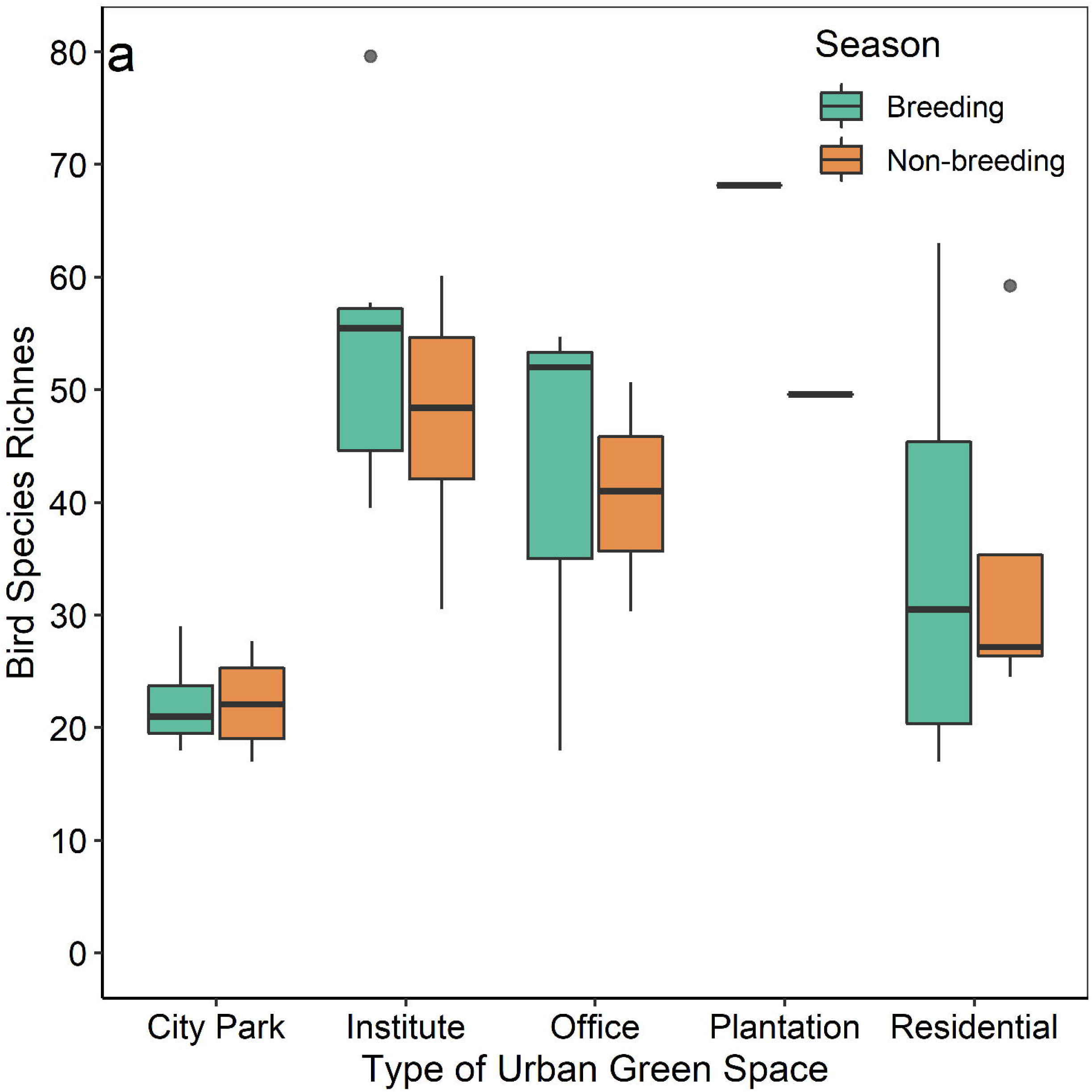
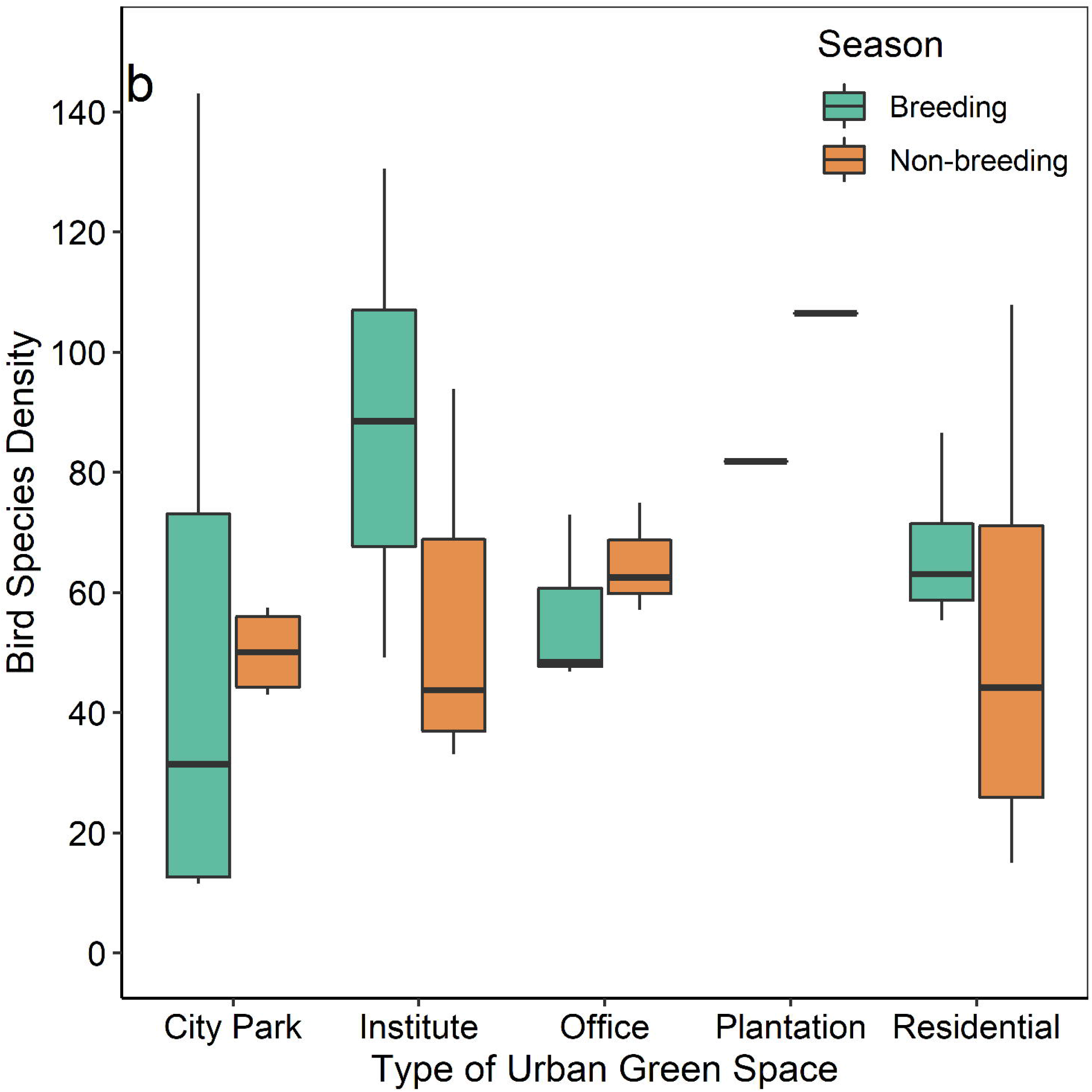
a) Overall bird species richness and b) density across urban green space types during breeding and non-breeding season.

Of the models explaining variation in bird species richness, the model containing only the urban green spaces size best explained the data during breeding season and non-breeding season (Table 2). The top model for the species richness explained 99% and 96% of the variation in data during breeding and non-breeding season, respectively (see electronic supplementary material A). Moreover, the effect size was more pronounced for the breeding than the non-breeding season (Table 2, Figure 3). At the local scale, top two models containing tree richness and a combination of tree and shrub richness, representing overall plant species richness, explained the data across breeding and non-breeding seasons. Two models cumulatively explained 98% of the variation in the data.

**Figure 3:**
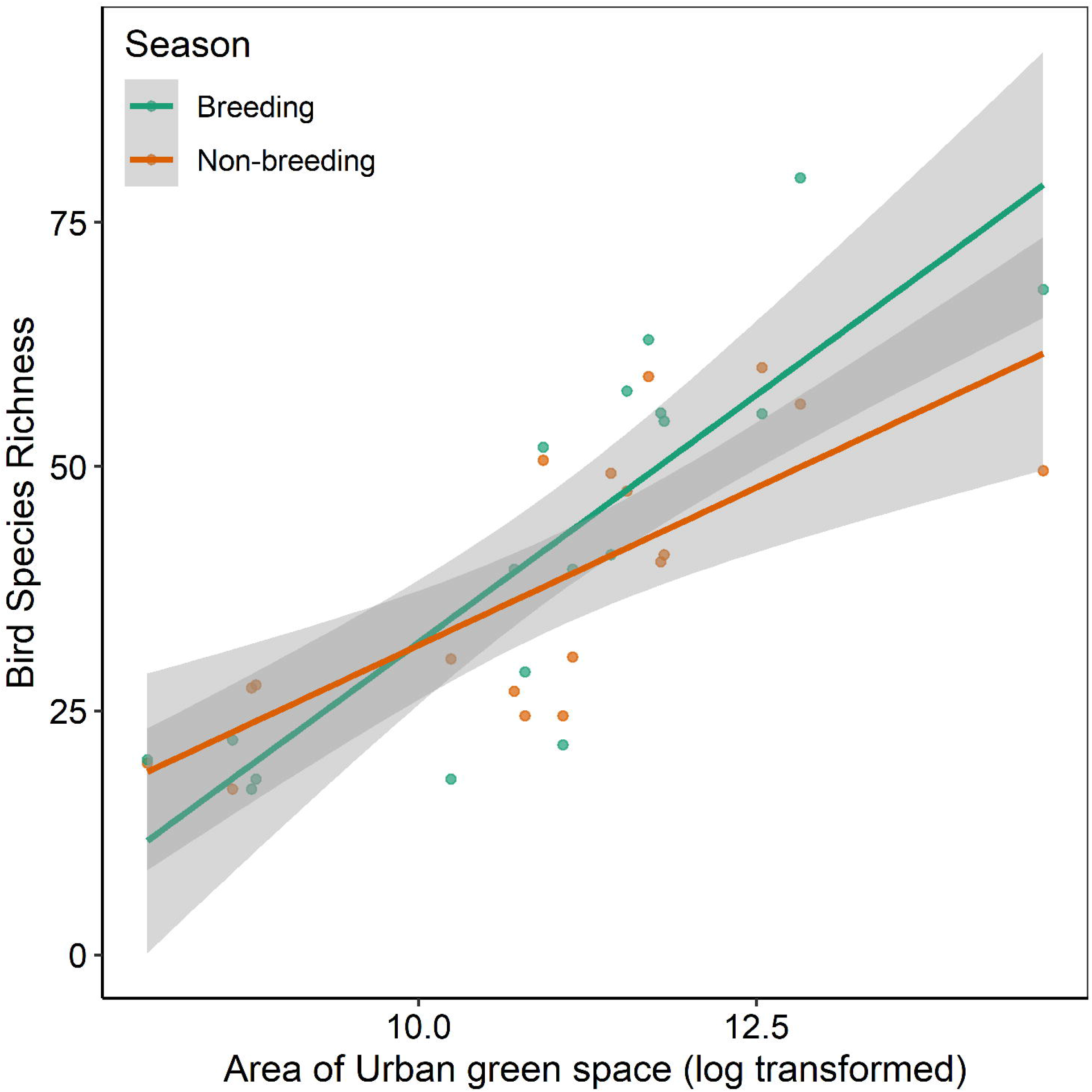
Relationship between bird species richness and area of the urban green spaces across breeding and non-breeding season.

**Table 2:**
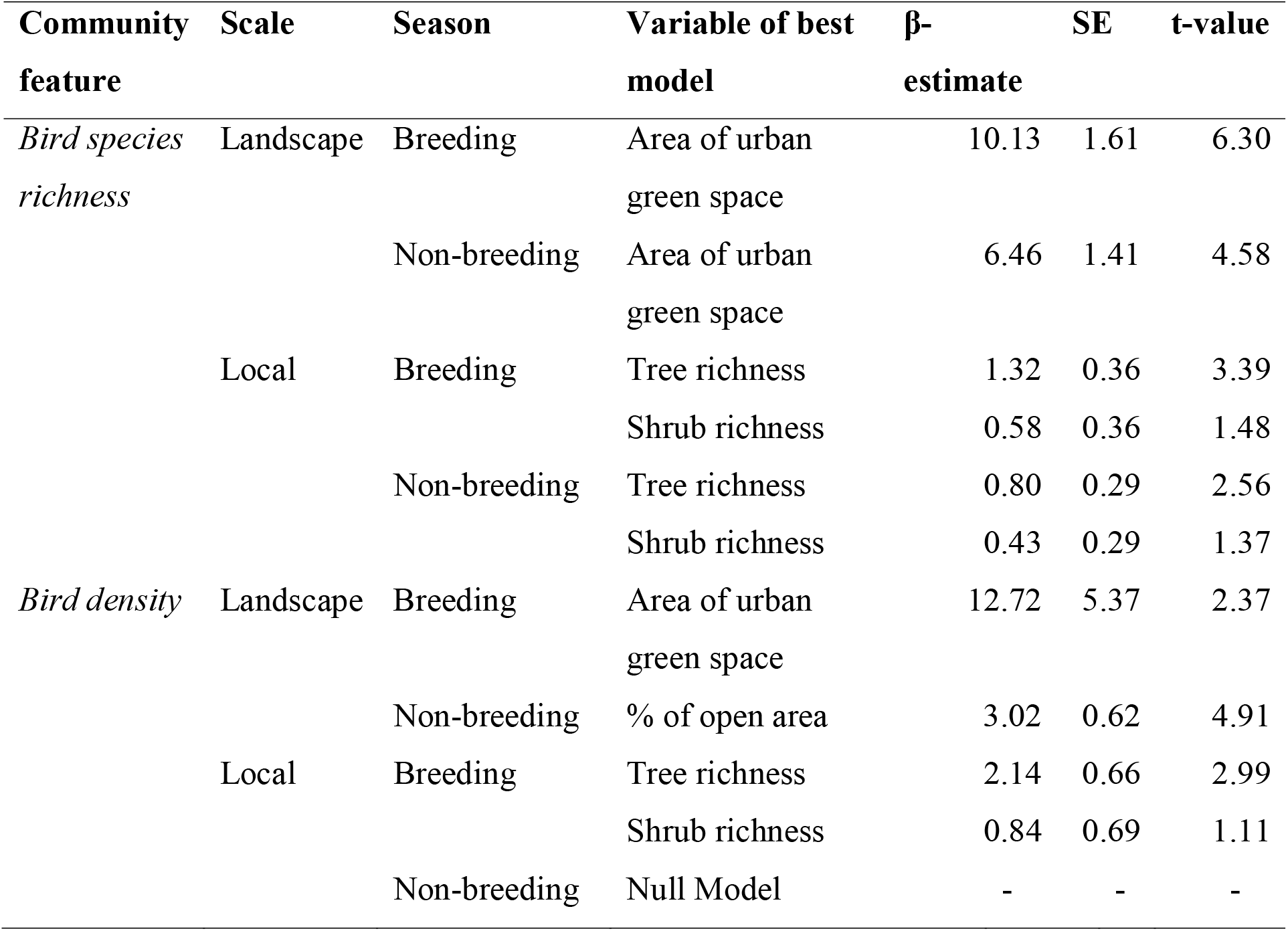
Summary of the best model showing variables, coefficient estimates, standard error, and associated t-value for effect of landscape and local scale features on bird community features during breeding and non-breeding season.

Overall bird density was explained by the park size during breeding (Table 2, Figure 4a) and at the local scale by additive effect of tree and shrub richness within the park (Table 2, Figure 4b & 4c). During non-breeding season, landscape level variable, i.e., percentage of open area in the matrix explained the variation in overall density (Table 2, Figure 4d). However, none of the local variables explained variation in density during non-breeding season (Table 2). The top model containing landscape level variables explained 50% and 88% of the variation in the overall density during breeding and non-breeding season respectively (see electronic supplementary material B). At the local scale the top model explained 94% of the variation in the overall bird density during the breeding season and no model was selected during the non-breeding season (see electronic supplementary material B).

**Figure 4:**
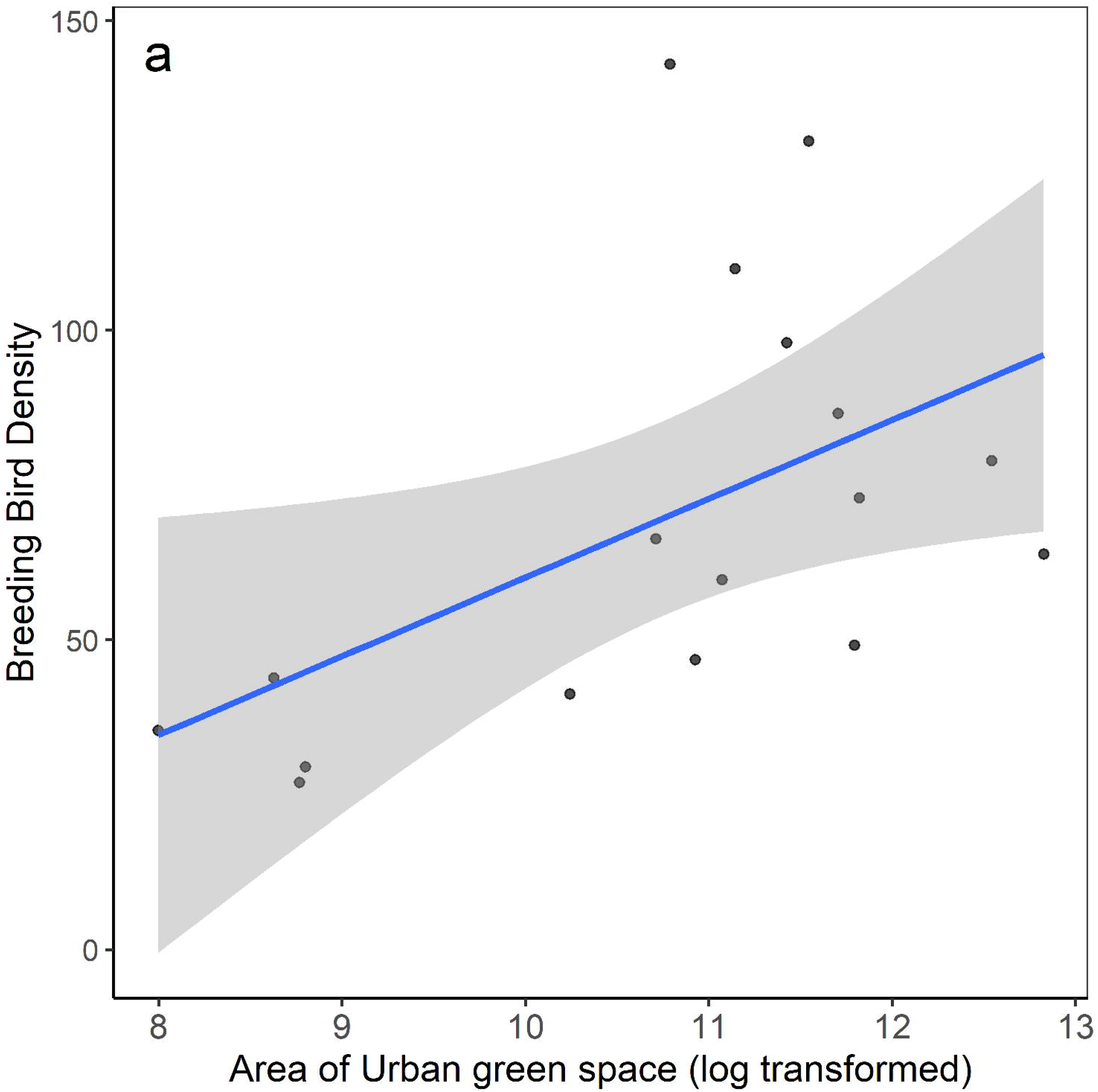
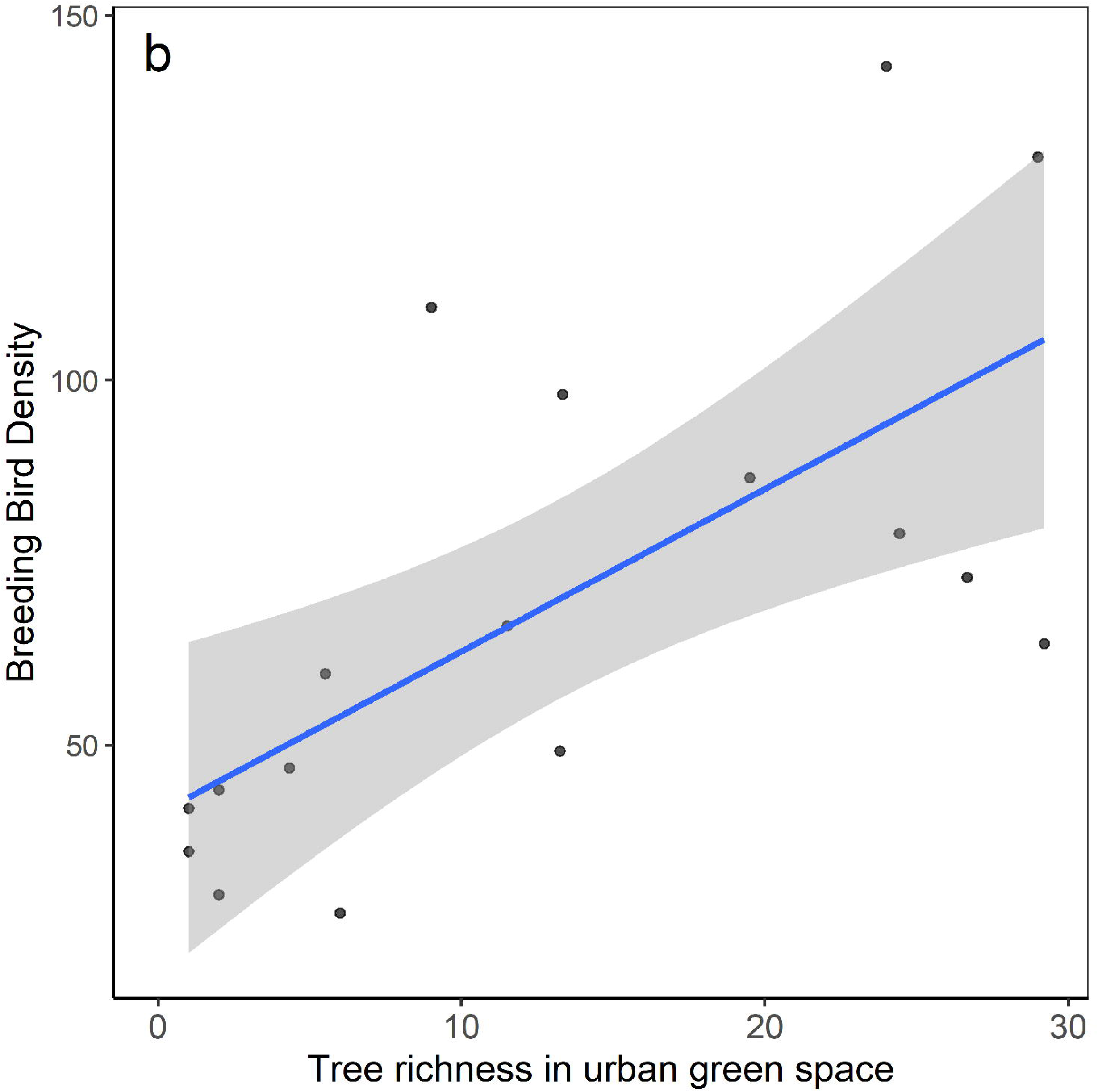
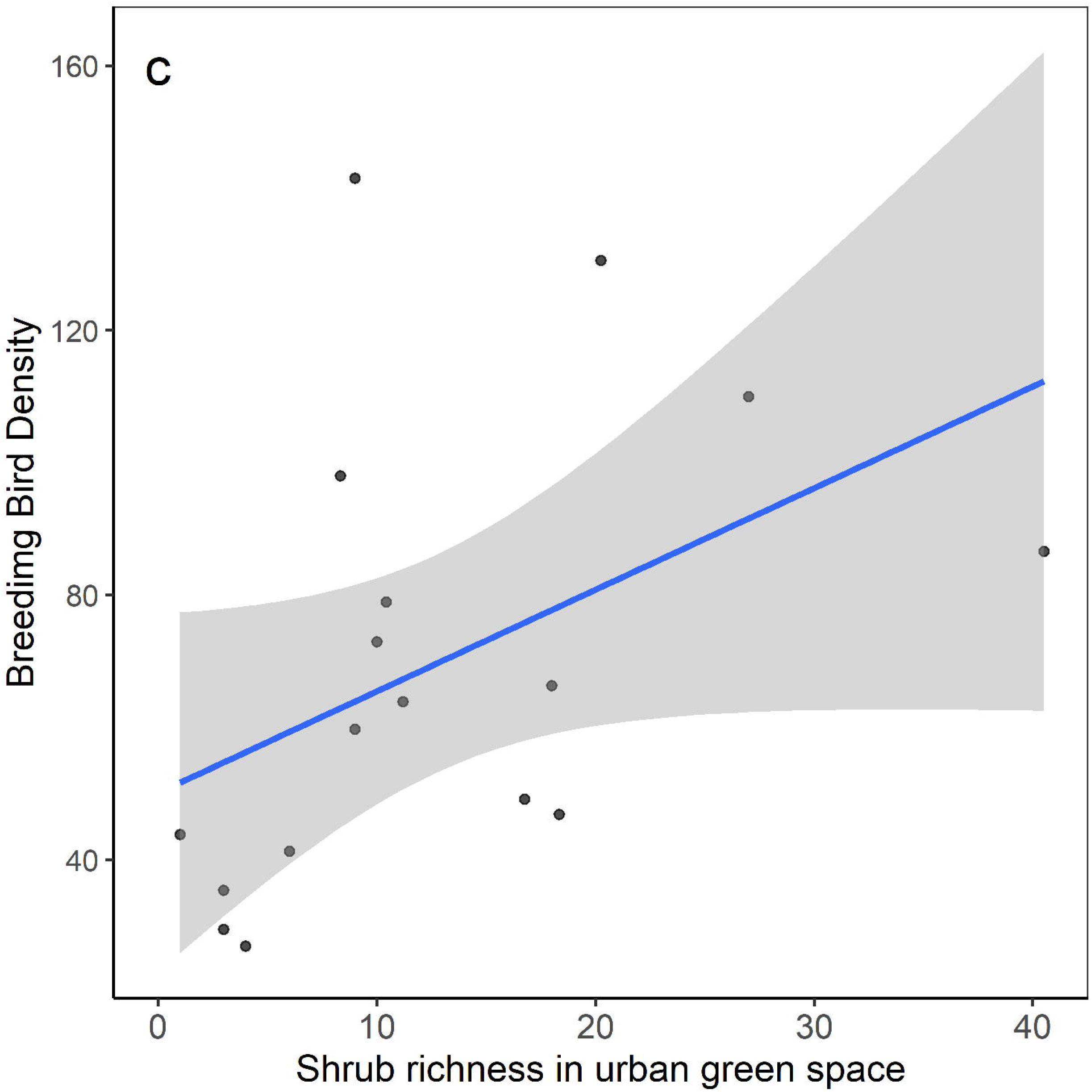
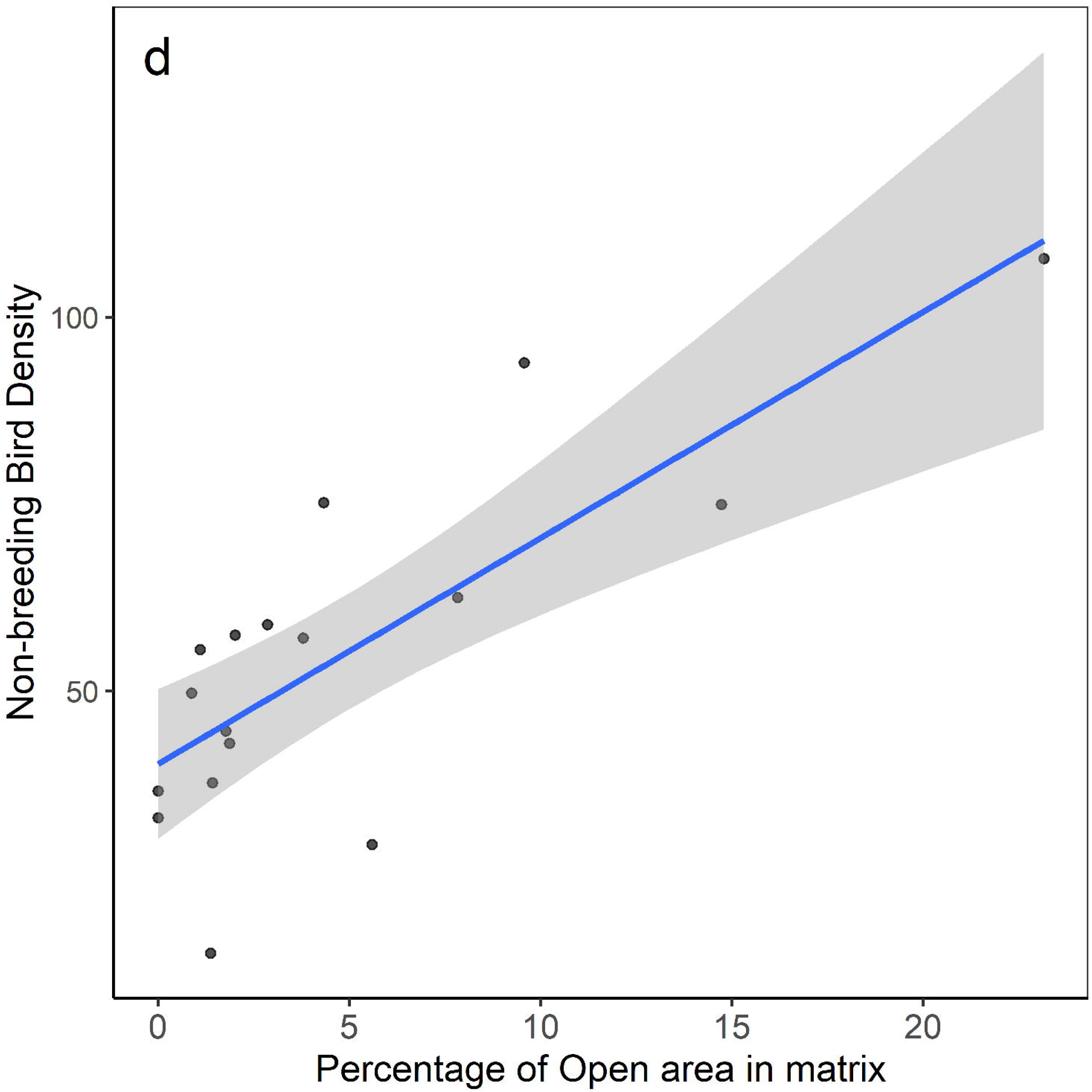
Relationship of overall bird density with parameters of the best models a), b) & c) for breeding and d) non-breeding season.

Richness of all insectivore guilds except ground insectivore increased with increasing area of the urban green spaces across breeding and non-breeding season (Table 3 & Table 4). Percentage of barren area in surrounding matrix caused increase in richness of sallying insectivore and granivore guild during breeding season. However, during non-breeding season only ground insectivore guild richness increased with increasing percentage of barren area in the surrounding matrix. Increase in percentage of built-up area in the matrix caused decline in richness of ground insectivore guild (Table 4). Frugivore-insectivore guild’s richness during non-breeding season increased with increasing percentage of agriculture area in the matrix. At local-scale tree species richness positively influenced richness of insectivorous guild richness during breeding and non-breeding season (Table 3 & 4). Richness of few guilds such as nectar-insectivore, fruit-seed-nectar and fruit-seed-nectar-insectivore was not explained by either landscape or local-scale variables (see electronic supplementary material C & D).

**Table 3:** Variable estimates, standard errors, and Z-value of the predictor variables of the best models results, for fine-foraging guild species richness during breeding season at 18 urban green spaces in Dehradun, India. * Model built without one extremely disturbed urban green space site. Only guilds for which removal resulted a change in best model is depicted here in addition to the analysis with all sites.

| **Fine-foraging Guild** | **Variable of best model** | **β-estimate** | **SE** | **Z-value** |
| --- | --- | --- | --- | --- |
| *Landscape scale* |  |  |  |  |
| Understory insectivore | Area of urban green space | 1.21 | 1.06 | 3.11 |
| Sallying insectivore | Area of urban green space | 1.28 | 1.11 | 2.12 |
|  | % of barren | 1.02 | 1.01 | 2.06 |
| Canopy insectivore | Area of urban green space | 1.47 | 1.11 | 3.60 |
| Ground insectivore | Null model | – | – | – |
| Frugivore insectivore | % of barren | -1.03 | 1.02 | -1.52 |
|  | Area of urban green space | 1.19 | 1.09 | 1.83 |
|  | % of agriculture | 1.03 | 1.03 | 1.15 |
| Trunk-bark foragers | Null Model | – | – | – |
| Trunk-bark foragers* | Area of urban green space | 1.82 | 1.34 | 2.03 |
| Granivore | % of barren | 1.02 | 1.01 | 2.08 |
| Granivore | % of agriculture | 1.05 | 1.03 | 1.28 |
| Omnivore | Area of urban green space | 1.27 | 1.09 | 2.63 |
| Nectar insectivore | Null model | – | – | – |
| Fruit-seed-nectar | Null model | – | – | – |
| Fruit-seed-nectar-insectivore | Null model | – | – | – |
| *Local scale* |  |  |  |  |
| Understory insectivore | Tree richness | 1.06 | 1.01 | 3.28 |
| Sallying insectivore | Tree richness | 1.03 | 1.01 | 2.15 |
|  | Tree girth | -1.01 | 1.00 | -1.77 |
| Canopy insectivore | Tree richness | 1.03 | 1.01 | 2.20 |
|  | Shrub cover | 1.21 | 1.06 | 2.78 |
|  | Shrub richness | 1.02 | 1.02 | 1.08 |
| Ground insectivore | Average tree height | -1.17 | 1.06 | -2.58 |
| Frugivore insectivore | Null Model | – | – | – |
| Trunk-bark foragers | Null Model | – | – | – |
| Trunk-bark foragers* | Tree richness | 1.05 | 1.02 | 2.10 |
| Granivore | Null model | – | – | – |
| Omnivore | Null model | – | – | – |
| Nectar insectivore | Null model | – | – | – |
| Fruit-seed-nectar | Null model | – | – | – |
| Fruit-seed-nectar-insectivore | Null model | – | – | – |

**Table 4:**
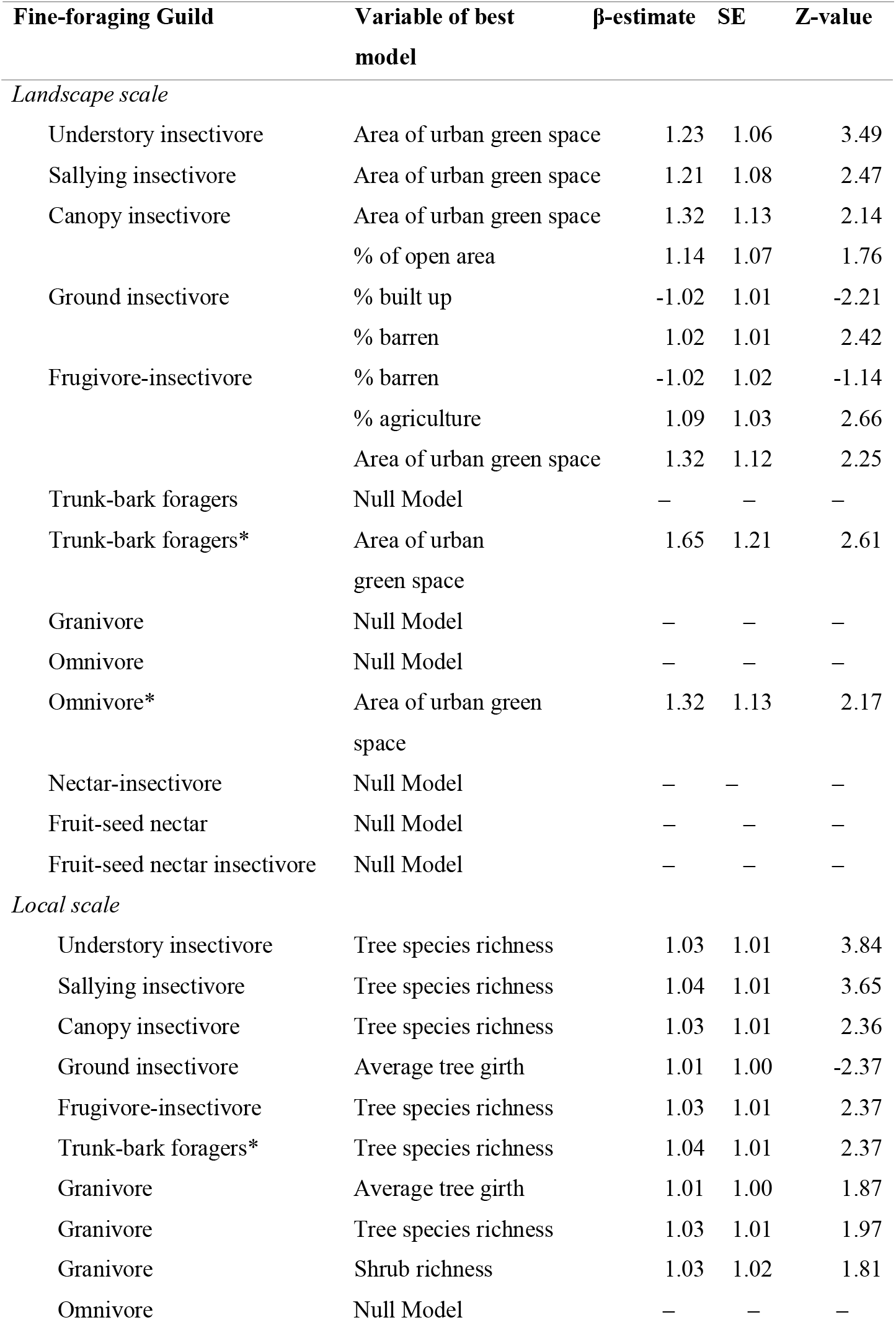
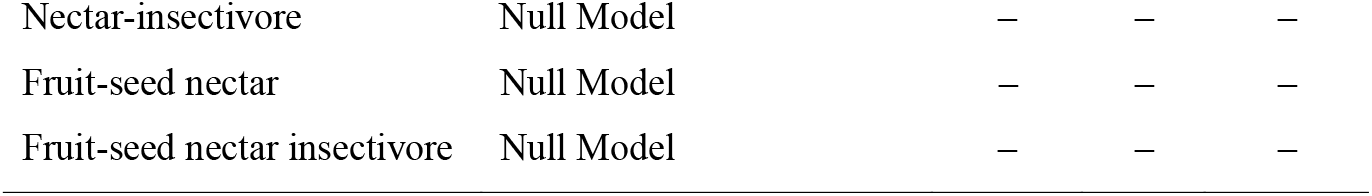
Variable estimates, standard errors, and Z-value of the predictor variables of the best models results, which predicted fine-foraging guild species richness during non-breeding season at 18 urban green spaces in Dehradun, India. * Model built without one extremely disturbed urban green space site. Only guilds for which removal resulted a change in best model is depicted here in addition to the analysis with all sites.

| **Fine-foraging Guild** | **Variable of best model** | **β-estimate** | **SE** | **Z-value** |
| --- | --- | --- | --- | --- |
| *Landscape scale* |  |  |  |  |
| Understory insectivore | Area of urban green space | 1.23 | 1.06 | 3.49 |
| Sallying insectivore | Area of urban green space | 1.21 | 1.08 | 2.47 |
| Canopy insectivore | Area of urban green space | 1.32 | 1.13 | 2.14 |
|  | % of open area | 1.14 | 1.07 | 1.76 |
| Ground insectivore | % built up | -1.02 | 1.01 | -2.21 |
|  | % barren | 1.02 | 1.01 | 2.42 |
| Frugivore-insectivore | % barren | -1.02 | 1.02 | -1.14 |
|  | % agriculture | 1.09 | 1.03 | 2.66 |
|  | Area of urban green space | 1.32 | 1.12 | 2.25 |
| Trunk-bark foragers | Null Model | — | — | — |
| Trunk-bark foragers* | Area of urban green space | 1.65 | 1.21 | 2.61 |
| Granivore | Null Model | — | — | — |
| Omnivore | Null Model | — | — | — |
| Omnivore* | Area of urban green space | 1.32 | 1.13 | 2.17 |
| Nectar-insectivore | Null Model | — | — | — |
| Fruit-seed nectar | Null Model | — | — | — |
| Fruit-seed nectar insectivore | Null Model | — | — | — |
| *Local scale* |  |  |  |  |
| Understory insectivore | Tree species richness | 1.03 | 1.01 | 3.84 |
| Sallying insectivore | Tree species richness | 1.04 | 1.01 | 3.65 |
| Canopy insectivore | Tree species richness | 1.03 | 1.01 | 2.36 |
| Ground insectivore | Average tree girth | 1.01 | 1.00 | -2.37 |
| Frugivore-insectivore | Tree species richness | 1.03 | 1.01 | 2.37 |
| Trunk-bark foragers* | Tree species richness | 1.04 | 1.01 | 2.37 |
| Granivore | Average tree girth | 1.01 | 1.00 | 1.87 |
| Granivore | Tree species richness | 1.03 | 1.01 | 1.97 |
| Granivore | Shrub richness | 1.03 | 1.02 | 1.81 |
| Omnivore | Null Model | — | — | — |
| Nectar-insectivore | Null Model | – | – | – |
| Fruit-seed nectar | Null Model | – | – | – |
| Fruit-seed nectar insectivore | Null Model | – | – | – |

### Bird species composition

Bird community composition in this study varied with urban green space area as well as with its type. As the urban green space become smaller in size, they become more dissimilar in bird species composition both during breeding (r= 0.32, p=0.001) and non-breeding (r= 0.32, p=0.005) season. Bird species composition also varied between urban green space types for breeding (r= 0.40, p=0.01) and non-breeding season (r= 0.56, p=0.001). We choose two dimensional NMDS because its correlation with the original data was only slightly lower than for a three-dimensional solution (*breeding season*: Linear fit R^2^ = 0.92 vs. 0.95; *non-breeding season*: Linear fit R^2^ = 0.88 vs. 0.92), while being easier to interpret. Overall goodness-of-fit calculated as *stress* of the solution was low across seasons (*breeding season*: Stress=0.11; *non-breeding season*: Stress=0.14). Spread of urban green spaces followed a similar pattern across seasons where large and medium sized urban green spaces clustered together but small-sized urban green spaces clustered in opposite direction (Figure 5a and 5b). Yet, there were a few sites that fell between the two clusters. Although geographically apart, large urban green spaces clustered very closely to each other whereas medium and small-sized urban green spaces showed huge variation in their bird composition. Landscape and local scale habitat parameters in this study significantly correlated with the NMDS axes. Interestingly some habitat parameters i.e., tree species richness, percentage of barren area, percentage of built-up and percentage of water, caused the differences in species composition across seasons. Whereas park size and percentage of agriculture land in the matrix influenced the community composition only during breeding season and average tree girth during non-breeding season. NMDS 1 strongly positively correlated with urban green space size, percentage of agricultural areas in the matrix and tree richness during breeding season aligning with large sized urban green spaces. In both seasons small urban green spaces aligned along a gradient of percentage of built-up in opposite direction to large and medium sized urban green spaces (Figure 5a and 5b).

**Figure 5:**
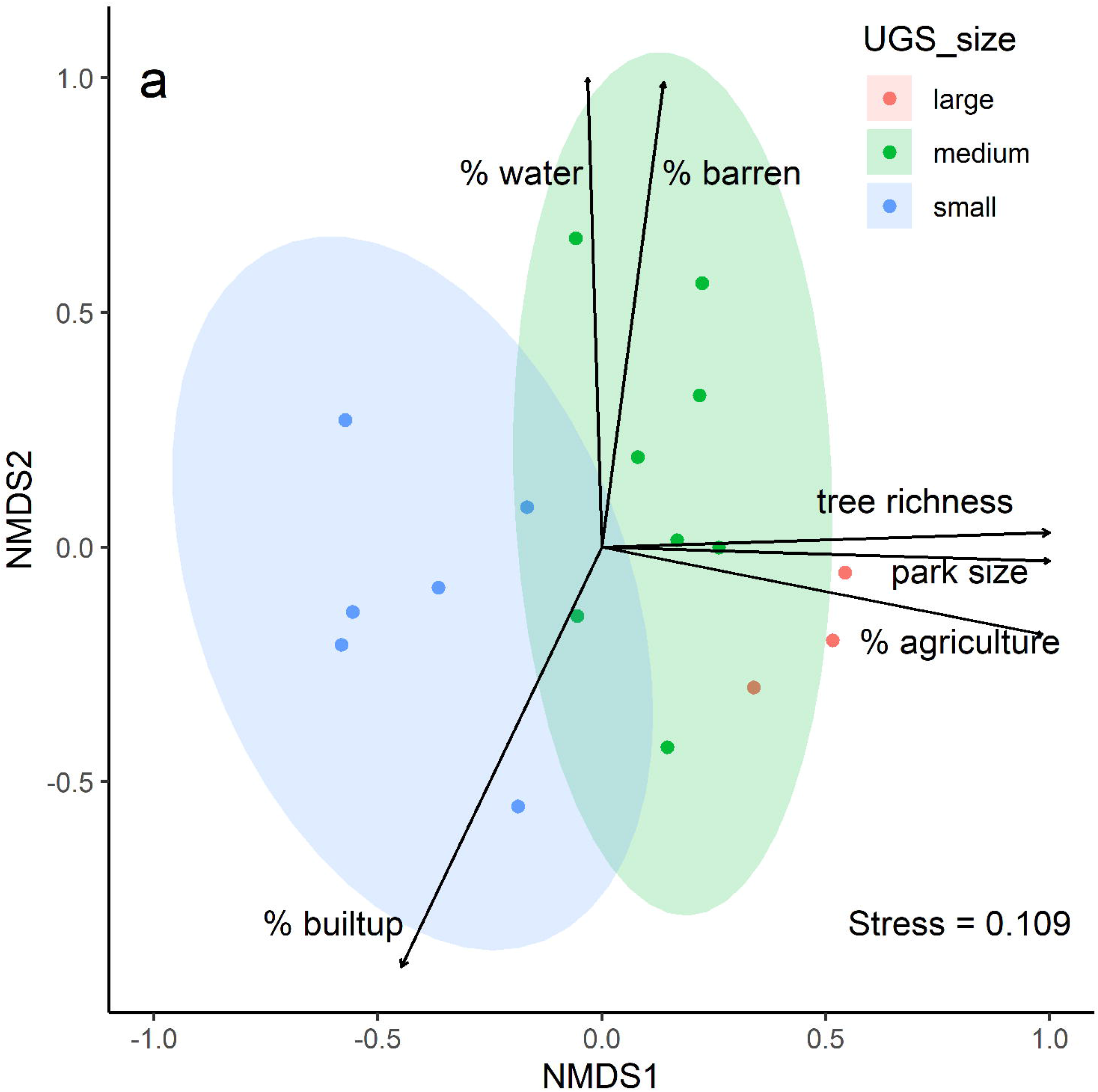
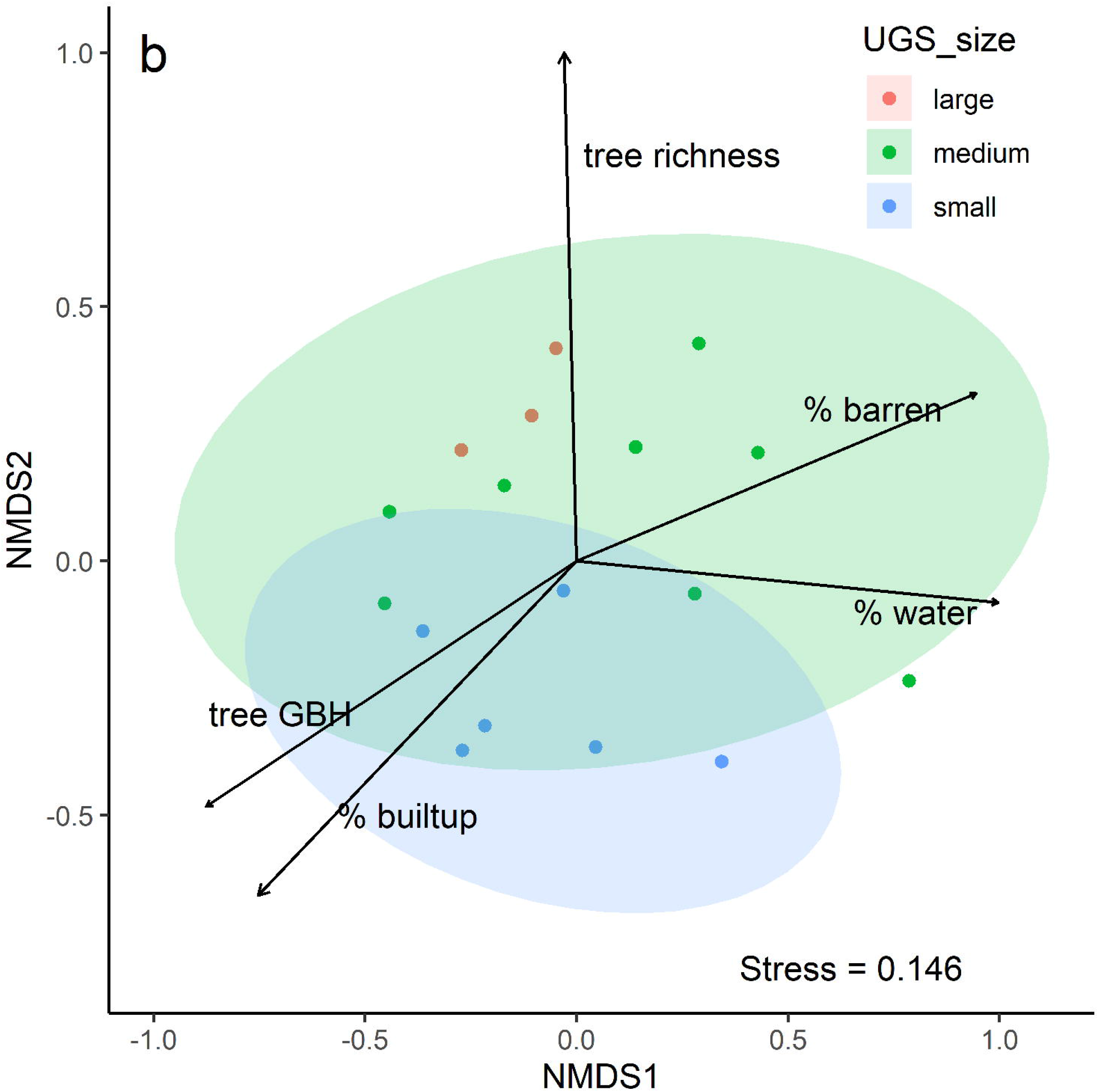
Nonmetric multidimensional scaling (NMDS) ordination of the bird community during a) breeding and b) non-breeding season at the 18 urban green space season in Dehradun, Uttarakhand. Plots represents sites according to their similarity in species composition. The arrows are vectors of habitat parameters arrows represent vectors of the significant factors that contributed to the ordination (p□<□0.05).

## Discussion

With the expansion of urbanization, it is becoming urgent to create and maintain spaces for urban biodiversity. Most importantly, such decision for planning and development of urban green spaces need to have its foundation in scientific knowledge. Information on urban green space features that improves their biodiversity potential has accumulated over the past few decades (Callaghan et al., 2018; Nielsen et al., 2014; Threlfall et al., 2017). Yet our knowledge is lopsided due to paucity of information from megadiverse developing countries (Callaghan et al., 2018). This study is the first attempt in the Himalayan state of Uttarakhand, northern India, to investigate the role of landscape and local scale variables in improving overall and specialist guild richness.

In consensus with the previous studies, our findings establish the value of landscape as well as local scale variables in influencing the bird species richness in urban green spaces(Callaghan et al., 2018; Dale, 2018; Mayorga et al., 2020). We found that urban green space size plays an overwhelmingly important role in supporting higher overall bird richness, density, and richness of specialized foraging guilds. A more encouraging result of this study is the significant role of tree and shrub richness at local scale for the breeding and non-breeding bird community (Table 2).

### Landscape scale determinants bird community characteristics

Species-area effect has been observed in studies conducted within urban green spaces of a single city and across cities as well. Callaghan et.al (2018) used citizen science data on bird observations from 112 urban green spaces spread across 51 cities and observed a significantly positive association between bird species richness of both terrestrial and water birds. Larger urban green spaces are expected to have diverse habitat providing foraging and nesting resources to a diversity of bird species (Matthies et al., 2017). Habitat heterogeneity or patchiness could provide safe refuges to birds for evading predation consequently leading to higher richness over long term (Willson et al., 2001). Although we did not quantify habitat diversity within urban green spaces but larger urban green spaces in this study had variety of habitats starting from regenerating forest areas, grasslands, scrubs, and vacant lots.

Another mechanism for larger urban green spaces to support higher bird richness is through increased within patch structural heterogeneity, a property of rich plant community. In this study we too observed a strong correlation between tree (r = 0.80, p < 0.001) and shrub richness (r = 0.55, p = 0.02) with the urban green space size. Larger urban green spaces with higher forage and nesting resources would have a direct effect on the abundance of the individual species. We also observed this effect of size on overall bird density during breeding season when the two imminent requirement of the bird are food and suitable nest site.

Overall bird density in this study increased with urban green space size during breeding season with percentage of open area in the matrix during non-breeding season (Figure 4a & 4d). This effect of park size on breeding bird abundance have been reported by other studies as well (Amaya-Espinel et al., 2019; Leveau & Leveau, 2016; Mayorga et al., 2020).

Linear relationship between urban green space size with breeding bird density could be attributed to productivity that is higher in green versus gray spaces (Shochat et al., 2006). Urban green spaces are also characterized by increased availability of subsided food, lower diversity and density of natural predators, prolonged breeding period of birds due to lack of seasonality subsequently leading to higher abundance of birds, especially urban exploiters, and adapters. Studies conducted in urban areas usually find the density of few urban exploiters contributing to this overall increase. In our study too, during breeding season 14% of species (17out of 123 recorded) contributed to 67%and 63% (14 out of 103 recorded) of the total bird abundance during breeding and non-breeding season, respectively. All these highly abundant (*Acridotheres tristis, Columba livia, Spilopelia chinensis, Orthotomus sutorius, Corvus splendens, Pycnonotus cafer* etc.) species were also characterized by widespread presence in majority of the sites. During non-breeding season, overall bird density in this study increased with increasing percentage of open area in the matrix, a land use with no or minimal management of vegetation. Non-breeding season in this area is marked by harsh winters and influx of 80% of the Himalayan birds to foothills and plains, avoiding even harsher winters in their breeding grounds. On arrival winter migrants in this region form mixed-foraging flocks with resident birds and show strong heterospecific attraction (Kaushik et al., 2012). These migrants often affects low and medium intensity agricultural fields than primary forest (Elsen et al., 2017).

Smaller urban green spaces in this study were nestled within the highly urbanized matrix, characterized by percentage of built-up (see Figure 5a and 5b). Although we did not find an impact of built-up area on the overall bird species richness but we did find a significant association with the bird species composition. Built-up area acts as barrier for movement between urban green spaces especially for disturbance sensitive ground dwelling and dispersal limited species. We indeed observed a decline in ground insectivore guild richness with increasing built-up cover in the matrix (Table 4). Although the study area is urbanizing at a fast rate, the presence of reserve forests around the boundary, remnant agricultural areas, old institutes with ample green cover, practices of home gardening seems to compensate for the effect of sealed area.

### Local scale determinants bird community characteristics

Tree richness at local scale had positive relationship with overall bird species richness, richness of most of the fine-foraging guilds, overall density, and bird species composition in our study. This relationship is also observed by other studies (da Silva et al., 2020; de Toledo et al., 2012; Khera et al., 2009). Increasing tree richness results in increase food and nesting resources for bird species. Tree richness is also positively related to foliage height diversity (Daniels et al. 1992) and therefore provide different foraging niches to the birds (MacArthur & MacArthur, 1961).

Contrary to our expectations we did not find effect of shrub richness on understory insectivore. We believe that this lack of relationship is due to the frequent control and management of shrub layer in the urban green spaces especially during the monsoon season to get rid of the insect and other pests. Effect of tree richness were more pronounced on the fine-foraging guilds of the birds and the species composition in our study. Richness of insectivores birds foraging in all stratum (understory, canopy, trunk-bark, air) increased with increasing tree richness. Tree richness potentially influence the richness of insectivorous guild by 1) increasing the foliage height diversity, 2) providing diverse food resources and by 3) providing cover from the predators (Evans et al., 2009). Other than tree richness, disturbance could negatively influence this group especially specialist group of Trunk-bark forager including woodpeckers. The largest site in this study was an old abandoned tea plantation with high native trees richness but the trees are heavily used for collecting firewood and fodder leading to lower richness of this specialized guild.

### Management implications

Our study provides further support for the park size as an important factor for conserving larger part of the bird diversity in urban areas. This finding is relevant for the city planners during planning stage as large urban green spaces can support a much larger array of bird species than the small ones. Additionally, green spaces within university campuses, offices, residential complex can further support the urban bird diversity. Although urban sprawl is expected to reduce the amount of barren and open areas but certain features of these land uses such as low or no management of shrubs could be incorporated in one portion of the urban green space. Another important finding of this study was the overwhelming role of the tree richness in improving the bird community characteristics at guild and community level. This finding could be used to plan improve the habitat quality of the small and medium parks for improving their conservation potential for bird community. Considering the lack of space for planning large urban green spaces within already planned cities focus should be on increasing native tree and shrub cover to imperiled protect ground insectivore guild.

## Acknowledgments

We thank the Rufford foundation for providing the small grant (Grant ID: 26698-1) to fund this research. We also thank Dr. V B Mathur, Director (Wildlife Institute of India), Dr. Geetika Mathur (Model School for visually handicapped), Saurabh Shanu (University of Petroleum and Energy Studies), Director General (Uttarakhand State Council for Science and Technology (Vigyan Dham), Archana Bahuguna, (Zoological Survey of India), staff of Tourism Office, Indu Singh, (Mahadevi Kanya Pathshala), for facilitating data collection in their institution’s premises. We are thankful to Oindrila Basu, Aashu tomar, Sipu kumar, Shilky, Rachna Yadav, Mangal Singh Bisht, Ananthu P, Jason B. Coutinho, Partho Sarathi, and Saloni Singh for their voluntary help for assisting ST and KM in data collection. We are immensely grateful to Inam Ali for assisting us in bird and vegetation sampling.

