## Appendix A for "Habitat patch size and tree species richness shape the bird community in urban green spaces of rapidly urbanizing region of India"

Table A: Model description, scale, degrees of freedom (df) Akaike Information Criteria (AICc), relative support for hypothesis (ΔAICc), of candidate models explaining bird species richness as a function of landscape and local variables during breeding and non-breeding season.

|  | Scale | Model | df | | AICc | ΔAICc | weight |
| --- | --- | --- | --- | --- | --- | --- | --- |
| Breeding Season | *Landscape* | Area of urban green space | 3 | | 142.8 | 0 | 0.999 |
|  |  | % of agriculture | 3 | | 157.2 | 14.36 | 0.001 |
|  |  | % of green cover+% of agriculture | 4 | | 159.7 | 16.85 | 0 |
|  |  | variation in green cover | 3 | | 161.5 | 18.65 | 0 |
|  |  | Null Model | 2 | | 162.4 | 19.54 | 0 |
|  |  | % of open area | 3 | | 164 | 21.17 | 0 |
|  |  | % of built-up | 3 | | 164.5 | 21.67 | 0 |
|  |  | % of green cover | 3 | | 165.2 | 22.4 | 0 |
|  |  | variation in open area | 3 | | 165.3 | 22.43 | 0 |
|  | *Local* | Tree richness | 3 | | 152.4 | 0 | 0.547 |
|  |  | Tree richness + Shrub richness | 4 | | 152.9 | 0.48 | 0.43 |
|  |  | Shrub richness | 3 | | 160.1 | 7.63 | 0.012 |
|  |  | Null Model | 2 | | 162.4 | 9.93 | 0.004 |
|  |  | Average tree girth | 3 | | 162.8 | 10.39 | 0.003 |
|  |  | Shrub cover | 3 | | 164 | 11.54 | 0.002 |
|  |  | Tree crown cover | 3 | | 165 | 12.56 | 0.001 |
|  |  | Average tree height | 3 | | 165.2 | 12.74 | 0.001 |
|  |  | Average tree crown cover+ Average shrub cover | 4 | | 167 | 14.6 | 0 |
|  | *Landscape* | Area of urban green space | 3 | | 138.1 | 0 | 0.958 |
| Non-Breeding Season |  | % of agricultural area | 3 | | 145.9 | 7.79 | 0.019 |
|  |  | % of agricultural area+% of green cover | 4 | | 147.9 | 9.78 | 0.007 |
|  |  | % of open area | 3 | | 149.5 | 11.4 | 0.003 |
|  |  | variation in green cover | 3 | | 149.7 | 11.65 | 0.003 |
|  |  | Null Model | 2 | | 150.3 | 12.18 | 0.002 |
|  |  | % of built-up | 3 | | 150.5 | 12.42 | 0.002 |
|  |  | % of built-up+I(% of built-up^2) | 4 | | 151 | 12.88 | 0.002 |
|  |  | % of open area | 3 | | 151.1 | 13.06 | 0.001 |
|  |  | % barren | 3 | | 153 | 14.89 | 0.001 |
|  |  | % green cover | 3 | | 153 | 14.91 | 0.001 |
|  |  | variation in open area | 3 | | 153.1 | 15.04 | 0.001 |
|  | *Local* | Tree richness | 3 | | 144.7 | 0.00 | 0.475 |
|  |  | Tree richness + Shrub richness | 4 | | 145.6 | 0.89 | 0.304 |
|  |  | Average tree girth | 3 | | 147.5 | 2.79 | 0.118 |
|  |  | Shrub cover | 3 | | 148.5 | 3.75 | 0.073 |
|  |  | Tree crown cover | | 3 | 152.2 | 7.47 | 0.01 |
|  |  | Average tree height | 3 | | 152.7 | 8.01 | 0.009 |
|  |  | Average tree crown cover+ Average shrub cover | 3 | | 153.0 | 8.31 | 0.007 |
