## Appendix B for "Habitat patch size and tree species richness shape the bird community in urban green spaces of rapidly urbanizing region of India"

Table B: Model description, scale, degrees of freedom (df) Akaike Information Criteria (AICc), relative support for hypothesis (ΔAICc), of candidate models explaining bird species density as a function of landscape and local variables during breeding and non-breeding season.

| Season | Scale | Model | df | AICc | ΔAICc | weight |
| --- | --- | --- | --- | --- | --- | --- |
| Breeding Season | *Landscape* | Area of urban green space | 3 | 170.1 | 0 | 0.50 |
|  |  | Null Model | 2 | 172.5 | 2.42 | 0.15 |
|  |  | % water | 3 | 174.9 | 4.83 | 0.05 |
|  |  | % builtup | 3 | 175 | 4.91 | 0.04 |
|  |  | % barren | 3 | 175.2 | 5.1 | 0.04 |
|  |  | % of built-up^2 | 3 | 175.3 | 5.21 | 0.04 |
|  |  | % of open area^2 | 3 | 175.3 | 5.27 | 0.04 |
|  |  | % of agricultural area | 3 | 175.3 | 5.27 | 0.04 |
|  |  | % of open area | 3 | 175.4 | 5.32 | 0.04 |
|  |  | % of green cover | 3 | 175.5 | 5.41 | 0.03 |
|  |  | % of green cover+ I (% of green cover^2) | 4 | 177.8 | 7.76 | 0.01 |
|  |  | % of green cover+% of water | 4 | 178.1 | 8 | 0.01 |
|  |  | % of built-up +% of green area | 4 | 178.4 | 8.38 | 0.01 |
|  |  | % of barren area+% builtup | 4 | 178.5 | 8.39 | 0.01 |
|  |  | % of green cover+ % agricultural area | 4 | 178.8 | 8.72 | 0.01 |
|  | *Local* | Tree richness | 3 | 173.9 | 0 | 0.68 |
|  |  | Tree richness+Shrub Richness | 4 | 175.8 | 1.89 | 0.26 |
|  |  | Shrub Richness | 3 | 180.7 | 6.78 | 0.02 |
|  |  | Null Model | 2 | 181.4 | 7.51 | 0.02 |
|  |  | Average Tree Height | 3 | 184.1 | 10.24 | 0.01 |
|  |  | Tree crown cover | 3 | 184.3 | 10.41 | 0.01 |
|  |  | Average Tree GBH | 3 | 184.3 | 10.41 | 0.01 |
|  |  | Shrub cover | 3 | 184.3 | 10.42 | 0.01 |
|  |  | Tree crown cover +Shrub cover | 4 | 187.7 | 13.77 | 0.01 |
| Non-Breeding Season | *Landscape* | % of open area | 3 | 146.1 | 0 | 0.88 |
|  |  | I(% of open area^2) | 3 | 150.2 | 4.07 | 0.16 |
|  |  | Null Model | 2 | 159.4 | 13.31 | 0.01 |
|  |  | % of agricultural area | 3 | 160.9 | 14.78 | 0.01 |
|  |  | % of built-up^2 | 3 | 161.1 | 15.02 | 0 |
|  |  | % water | 3 | 162.1 | 15.95 | 0 |
|  |  | % builtup | 3 | 162.1 | 15.98 | 0 |
|  |  | Area of urban green space | 3 | 162.1 | 15.98 | 0 |
|  |  | % of barren area+% builtup | 4 | 162.1 | 16.03 | 0 |
|  |  | % barren | 3 | 162.2 | 16.05 | 0 |
|  |  | % of green cover | 3 | 162.4 | 16.3 | 0 |
|  |  | % of green cover+ % agricultural area | 4 | 164.3 | 18.14 | 0 |
|  |  | % of green cover+% of water | 4 | 165.3 | 19.23 | 0 |
|  |  | % of built-up +% of green area | 4 | 165.5 | 19.41 | 0 |
|  |  | % of green cover+ I(% of green cover^2) | 4 | 165.8 | 19.72 | 0 |
|  | *Local* | Null Model | 3 | 173.9 | 0 | 0.68 |
|  |  | Tree richness+Shrub Richness | 4 | 175.8 | 1.89 | 0.26 |
|  |  | Tree richness | 3 | 180.7 | 6.78 | 0.02 |
|  |  | Shrub Richness | 2 | 181.4 | 7.51 | 0.02 |
|  |  | Tree crown cover | 3 | 184.1 | 10.24 | 0.01 |
|  |  | Shrub cover | 3 | 184.3 | 10.41 | 0.01 |
|  |  | Average Tree Height | 3 | 184.3 | 10.41 | 0.01 |
|  |  | Average Tree GBH | 3 | 184.3 | 10.42 | 0.01 |
|  |  | Tree crown cover +Shrub cover | 4 | 187.7 | 13.77 | 0.01 |
