## Appendix C for "Habitat patch size and tree species richness shape the bird community in urban green spaces of rapidly urbanizing region of India"

Table C: Model description, scale, degrees of freedom (df), Akaike Information Criteria (AICc), relative support for hypothesis (AICc), of candidate models explaining find-foraging guild richness as a function of landscape and local variables during non-breeding season.

| Guild | Scale | Model | df | AICc | ΔAICc |  | weight |
| --- | --- | --- | --- | --- | --- | --- | --- |
| Understory insectivore | *Landscape* | Area of urban green space | 2 | 80.4 | 0 |  | 0.902 |
|  |  | % barren+% agriculture | 3 | 85.3 | 4.99 |  | 0.075 |
|  |  | Null Model | 1 | 90 | 9.61 |  | 0.007 |
|  |  | % barren | 2 | 90.4 | 10.01 |  | 0.006 |
|  |  | % green | 2 | 91.6 | 11.19 |  | 0.003 |
|  |  | % builtup | 2 | 92.1 | 11.74 |  | 0.003 |
|  |  | % open | 2 | 92.5 | 12.16 |  | 0.002 |
|  |  | % barren+% open | 3 | 93.2 | 12.83 |  | 0.001 |
|  | *Local* | tree richness | 2 | 77.4 | 0 |  | 0.794 |
|  |  | tree richness+shrub richness | 3 | 80.3 | 2.91 |  | 0.185 |
|  |  | average tree girth | 2 | 85.4 | 8.03 |  | 0.014 |
|  |  | shrub richness | 2 | 89.5 | 12.1 |  | 0.002 |
|  |  | Null Model | 1 | 90 | 12.55 |  | 0.001 |
|  |  | tree crown cover | 2 | 91.2 | 13.79 |  | 0.001 |
|  |  | shrub cover | 2 | 91.3 | 13.87 |  | 0.001 |
|  |  | average tree height | 2 | 91.8 | 14.38 |  | 0.001 |
|  |  | tree crown cover+shrub cover | 3 | 92.9 | 15.44 |  | 0 |
| Sallying insectivores | *Landscape* | Area of urban green space | 2 | 70.6 | 0 |  | 0.629 |
|  |  | Null Model | 1 | 74.1 | 3.52 |  | 0.108 |
|  |  | % barren | 2 | 74.7 | 4.12 |  | 0.08 |
|  |  | % builtup | 2 | 75.4 | 4.8 |  | 0.057 |
|  |  | % open | 2 | 76.3 | 5.64 |  | 0.037 |
|  |  | % barren+ % agriculture | 3 | 76.3 | 5.65 |  | 0.037 |
|  |  | % green | 2 | 76.7 | 6.04 |  | 0.031 |
|  |  | % barren+% open | 3 | 77.5 | 6.91 |  | 0.02 |
|  | *Local* | tree richness | 2 | 62.7 | 0 |  | 0.756 |
|  |  | tree richness+shrub richness | 3 | 65.4 | 2.77 |  | 0.189 |
|  |  | average tree girth | 2 | 68.4 | 5.77 |  | 0.042 |
|  |  | shrub richness | 2 | 72.4 | 9.78 |  | 0.006 |
|  |  | Null Model | 1 | 74.1 | 11.46 |  | 0.002 |
|  |  | tree crown cover | 2 | 74.2 | 11.56 |  | 0.002 |
|  |  | average tree height | 2 | 75.3 | 12.65 |  | 0.001 |
|  |  | shrub cover | 2 | 76.7 | 14 |  | 0.001 |
|  |  | tree crown cover+shrub cover | 3 | 77.1 | 14.45 |  | 0.001 |
| Canopy insectivore | *Landscape* | Area of urban green space | 2 | 59.7 | 0 |  | 0.358 |
|  |  | % open | 2 | 59.8 | 0.17 |  | 0.329 |
|  |  | Null Model | 1 | 62.4 | 2.75 |  | 0.09 |
|  |  | % barren+% open | 3 | 62.6 | 2.89 |  | 0.084 |
|  |  | % builtup | 2 | 63.3 | 3.64 |  | 0.058 |
|  |  | % green | 2 | 64.2 | 4.5 |  | 0.038 |
|  |  | % barren | 2 | 65 | 5.3 |  | 0.025 |
|  |  | % barren+ % agriculture | 3 | 65.8 | 6.11 |  | 0.017 |
|  | *Local* | tree richness | 2 | 59.1 | 0 |  | 0.51 |
|  |  | tree richness+shrub richness | 3 | 61.8 | 2.68 |  | 0.134 |
|  |  | Null Model | 1 | 62.4 | 3.3 |  | 0.098 |
|  |  | shrub richness | 2 | 62.6 | 3.45 |  | 0.091 |
|  |  | average tree girth | 2 | 63.1 | 4 |  | 0.069 |
|  |  | tree crown cover | 2 | 64.7 | 5.59 |  | 0.031 |
|  |  | average tree height | 2 | 64.8 | 5.67 |  | 0.03 |
|  |  | shrub cover | 2 | 64.8 | 5.67 |  | 0.03 |
|  |  | tree crown cover+shrub cover | 3 | 67.4 | 8.31 |  | 0.008 |
| Ground insectivore | *Landscape* | % builtup | 2 | 65.4 | 0 |  | 0.352 |
|  |  | % barren | 2 | 65.7 | 0.21 |  | 0.317 |
|  |  | % barren+% open | 3 | 68.1 | 2.61 |  | 0.095 |
|  |  | % barren+ % agriculture | 3 | 68.5 | 3.06 |  | 0.076 |
|  |  | Null Model | 1 | 68.5 | 3.08 |  | 0.075 |
|  |  | Area of urban green space | 2 | 70.1 | 4.62 |  | 0.035 |
|  |  | % green | 2 | 70.5 | 5.05 |  | 0.028 |
|  |  | % open | 2 | 71.1 | 5.62 |  | 0.021 |
|  | *Local* | average tree girth | 2 | 64.3 | 0 |  | 0.549 |
|  |  | average tree height | 2 | 66.9 | 2.56 |  | 0.153 |
|  |  | tree richness | 2 | 67.7 | 3.39 |  | 0.101 |
|  |  | Null Model | 1 | 68.5 | 4.23 |  | 0.066 |
|  |  | tree crown cover | 2 | 69.1 | 4.77 |  | 0.051 |
|  |  | shrub richness | 2 | 70.5 | 6.16 |  | 0.025 |
|  |  | tree richness+shrub richness | 3 | 70.6 | 6.3 |  | 0.024 |
|  |  | shrub cover | 2 | 71 | 6.68 |  | 0.019 |
|  |  | tree crown cover+shrub cover | 3 | 71.8 | 7.55 |  | 0.013 |
| Granivore | Landscape | Null model | 1 | 55.2 | 0 |  | 0.236 |
|  |  | Area of urban green space | 2 | 55.4 | 0.28 |  | 0.206 |
|  |  | % barren | 2 | 55.7 | 0.5 |  | 0.184 |
|  |  | % builtup | 2 | 56.7 | 1.56 |  | 0.108 |
|  |  | % green | 2 | 57 | 1.83 |  | 0.094 |
|  |  | % open | 2 | 57.7 | 2.55 |  | 0.066 |
|  |  | % barren+ % agriculture | 9 3 | 57.9 | 2.78 |  | 0.059 |
|  |  | % barren+% open | 3 | 58.4 | 3.26 |  | 0.046 |
|  | *Local* | average tree girth | 2 | 52.8 | 0 |  | 0.289 |
|  |  | tree richness | 2 | 53.1 | 0.23 |  | 0.258 |
|  |  | shrub richness | 2 | 54.3 | 1.46 |  | 0.139 |
|  |  | tree richness+shrub richness | 3 | 55.1 | 2.25 |  | 0.094 |
|  |  | Null Model | 1 | 55.2 | 2.33 |  | 0.09 |
|  |  | tree crown cover | 2 | 56.4 | 3.6 |  | 0.048 |
|  |  | average tree height | 2 | 56.6 | 3.76 |  | 0.044 |
|  |  | shrub cover | 2 | 57.6 | 4.72 |  | 0.027 |
|  |  | tree crown cover+shrub cover | 3 | 59.2 | 6.33 |  | 0.012 |
| Nectar insectivore | *Landscape* | Null Model | 1 | 56.8 | 0 |  | 0.32 |
|  |  | Area of urban green space | 2 | 57.3 | 0.46 |  | 0.255 |
|  |  | % builtup | 2 | 59.3 | 2.52 |  | 0.091 |
|  |  | % open | 2 | 59.3 | 2.52 |  | 0.091 |
|  |  | % green | 2 | 59.4 | 2.54 |  | 0.09 |
|  |  | % barren | 2 | 59.4 | 2.54 |  | 0.09 |
|  |  | % barren+ % agriculture | 3 | 60.8 | 4.02 |  | 0.043 |
|  |  | % barren+% open | 3 | 62.3 | 5.43 |  | 0.021 |
|  | *Local* | Null Model | 1 | 56.8 | 0 |  | 0.28 |
|  |  | tree crown cover | 2 | 58.2 | 1.36 |  | 0.142 |
|  |  | shrub cover | 2 | 58.2 | 1.41 |  | 0.138 |
|  |  | tree richness | 2 | 58.8 | 1.93 |  | 0.107 |
|  |  | average tree height | 2 | 59.2 | 2.37 |  | 0.085 |
|  |  | average tree girth | 2 | 59.3 | 2.47 |  | 0.081 |
|  |  | shrub richness | 2 | 59.4 | 2.55 |  | 0.078 |
|  |  | tree crown cover+shrub cover | 3 | 59.9 | 3.07 |  | 0.06 |
|  |  | tree richness+shrub richness | 3 | 61.5 | 4.65 |  | 0.027 |
| fruit-seed nectar insectivore | *Landscape* | % green | 2 | 72.6 | 0 |  | 0.281 |
|  |  | Null Model | 1 | 73 | 0.33 |  | 0.238 |
|  |  | % builtup | 2 | 74 | 1.39 |  | 0.14 |
|  |  | % open | 2 | 74.1 | 1.49 |  | 0.133 |
|  |  | Area of urban green space | 2 | 75 | 2.39 |  | 0.085 |
|  |  | % barren | 2 | 75.5 | 2.87 |  | 0.067 |
|  |  | % barren+% open | 3 | 77 | 4.4 |  | 0.031 |
|  |  | % barren+ % agriculture | 3 | 77.6 | 4.94 |  | 0.024 |
|  | *Local* | tree richness | 2 | 72.2 | 0 |  | 0.306 |
|  |  | Null Model | 1 | 73 | 0.78 |  | 0.207 |
|  |  | tree richness + shrub richness | 3 | 74.3 | 2.09 |  | 0.108 |
|  |  | average tree girth | 2 | 74.6 | 2.39 |  | 0.093 |
|  |  | tree crown cover | 2 | 75.1 | 2.9 |  | 0.072 |
|  |  | shrub cover | 2 | 75.1 | 2.93 |  | 0.071 |
|  |  | average tree height | 2 | 75.2 | 3.05 |  | 0.067 |
|  |  | shrub richness | 2 | 75.5 | 3.33 |  | 0.058 |
|  |  | tree crown cover+shrub cover | 3 | 77.6 | 5.42 |  | 0.02 |
| Fruit-seed nectar | *Landscape* | Null Model | 1 | 48.9 | 0 |  | 0.288 |
|  |  | Area of urban green space | 2 | 49.5 | 0.62 |  | 0.211 |
|  |  | % barren | 2 | 50.7 | 1.79 |  | 0.118 |
|  |  | % builtup | 2 | 51.1 | 2.2 |  | 0.096 |
|  |  | % open | 2 | 51.2 | 2.3 |  | 0.091 |
|  |  | % green | 2 | 51.3 | 2.47 |  | 0.084 |
|  |  | % barren+ % agriculture | 3 | 51.4 | 2.57 |  | 0.079 |
|  |  | % barren+% open | 3 | 53.2 | 4.29 |  | 0.034 |
|  | *Local* | tree richness | 2 | 48 | 0 |  | 0.316 |
|  |  | Null Model | 1 | 48.9 | 0.83 |  | 0.208 |
|  |  | average tree height | 2 | 49.9 | 1.89 |  | 0.123 |
|  |  | tree richness+shrub richness | 3 | 50.9 | 2.87 |  | 0.075 |
|  |  | shrub richness | 2 | 51 | 2.96 |  | 0.072 |
|  |  | shrub cover | 2 | 51.1 | 3.07 |  | 0.068 |
|  |  | tree crown cover | 2 | 51.3 | 3.25 |  | 0.062 |
|  |  | average tree girth | 2 | 51.4 | 3.37 |  | 0.059 |
|  |  | tree crown cover+shrub cover | 3 | 53.9 | 5.84 |  | 0.017 |
| Raptor | *Landscape* | Null Model | 1 | 48.3 | 0 |  | 0.329 |
|  |  | Area of urban green space | 2 | 49.7 | 1.31 |  | 0.171 |
|  |  | % builtup | 2 | 50.3 | 1.95 |  | 0.124 |
|  |  | % barren | 2 | 50.4 | 2.09 |  | 0.116 |
|  |  | % open | 2 | 50.7 | 2.34 |  | 0.102 |
|  |  | % green | 2 | 50.9 | 2.52 |  | 0.093 |
|  |  | % barren+ % agriculture | 3 | 52.7 | 4.36 |  | 0.037 |
|  |  | % barren+% open | 3 | 53.2 | 4.88 |  | 0.029 |
|  | *Local* | Null Model | 1 | 48.3 | 0 |  | 0.324 |
|  |  | tree richness | 2 | 50.5 | 2.14 |  | 0.111 |
|  |  | tree crown cover | 2 | 50.5 | 2.19 |  | 0.108 |
|  |  | average tree girth | 2 | 50.6 | 2.25 |  | 0.105 |
|  |  | average tree height | 2 | 50.6 | 2.27 |  | 0.104 |
|  |  | shrub cover | 2 | 50.7 | 2.36 |  | 0.1 |
|  |  | shrub richness | 2 | 50.8 | 2.49 |  | 0.093 |
|  |  | tree crown cover+shrub cover | 3 | 53.2 | 4.9 |  | 0.028 |
|  |  | tree richness+shrub richness | 3 | 53.4 | 5.04 |  | 0.026 |
| Omnivore | *Landscape* | Area of urban green space | 2 | 63.2 | 0 |  | 0.441 |
|  |  | Null Model | 1 | 64.5 | 1.28 |  | 0.233 |
|  |  | % builtup | 2 | 66.6 | 3.43 |  | 0.079 |
|  |  | % green | 2 | 66.8 | 3.56 |  | 0.074 |
|  |  | % open | 2 | 67 | 3.78 |  | 0.066 |
|  |  | % barren | 2 | 67 | 3.82 |  | 0.065 |
|  |  | % barren+ % agriculture | 3 | 68.9 | 5.72 |  | 0.025 |
|  |  | % barren+% open | 3 | 69.9 | 6.68 |  | 0.016 |
|  | *Local* | tree richness | 2 | 62.6 | 0 |  | 0.398 |
|  |  | Null Model | 1 | 64.5 | 1.9 |  | 0.154 |
|  |  | tree richness+shrub richness | 3 | 65.4 | 2.84 |  | 0.096 |
|  |  | shrub richness | 2 | 65.5 | 2.94 |  | 0.092 |
|  |  | tree crown cover | 2 | 66.2 | 3.64 |  | 0.064 |
|  |  | average tree height | 2 | 66.3 | 3.73 |  | 0.062 |
|  |  | average tree girth | 2 | 66.5 | 3.87 |  | 0.057 |
|  |  | shrub cover | 2 | 66.5 | 3.92 |  | 0.056 |
|  |  | tree crown cover+shrub cover | 3 | 68.6 | 6 |  | 0.02 |
| Frugivore insectivore | Landscape | % barren+ % agriculture | 3 | 58.6 | 0 |  | 0.456 |
|  |  | Area of urban green space | 2 | 59.1 | 0.58 |  | 0.341 |
|  |  | Null Model | 1 | 62.4 | 3.88 |  | 0.065 |
|  |  | % barren | 2 | 63 | 4.42 |  | 0.05 |
|  |  | % builtup | 2 | 63.7 | 5.13 |  | 0.035 |
|  |  | % green | 2 | 64.7 | 6.1 |  | 0.022 |
|  |  | % open | 2 | 64.9 | 6.33 |  | 0.019 |
|  |  | % barren+% open | 3 | 65.9 | 7.32 |  | 0.012 |
|  | Local | tree richness | 2 | 58.7 | 0 |  | 0.58 |
|  |  | tree richness+shrub richness | 3 | 61.3 | 2.6 |  | 0.158 |
|  |  | Null Model | 1 | 62.4 | 3.75 |  | 0.089 |
|  |  | average tree height | 2 | 63.9 | 5.25 |  | 0.042 |
|  |  | average tree girth | 2 | 64.1 | 5.39 |  | 0.039 |
|  |  | shrub richness | 2 | 64.6 | 5.9 |  | 0.03 |
|  |  | tree crown cover | 2 | 64.7 | 6 |  | 0.029 |
|  |  | shrub cover | 2 | 65 | 6.29 |  | 0.025 |
|  |  | tree crown cover+shrub cover | 3 | 67.6 | 8.9 |  | 0.007 |
| Trunk-bark foragers | Landscape | Area of urban green space | 2 | 52.6 | 0 |  | 0.824 |
|  |  | Null model | 3 | 58.4 | 5.8 |  | 0.045 |
|  |  | % built-up | 1 | 58.7 | 6.04 |  | 0.04 |
|  |  | % barren | 2 | 58.8 | 6.18 |  | 0.038 |
|  |  | % barren+ % agriculture | 2 | 60.2 | 7.61 |  | 0.018 |
|  |  | % open | 2 | 61 | 8.35 |  | 0.013 |
|  |  | % green | 2 | 61 | 8.37 |  | 0.013 |
|  |  | % barren +% open | 3 | 61.7 | 9.07 |  | 0.009 |
|  | Local | tree richness | 2 | 57.2 | 0 |  | 0.559 |
|  |  | tree richness+shrub richness | 3 | 60.1 | 2.91 |  | 0.131 |
|  |  | Null Model | 1 | 60.6 | 3.37 |  | 0.104 |
|  |  | average tree height | 2 | 61.9 | 4.7 |  | 0.053 |
|  |  | shrub richness | 2 | 62.1 | 4.87 |  | 0.049 |
|  |  | shrub cover | 2 | 62.7 | 5.5 |  | 0.036 |
|  |  | tree crown cover | 2 | 63 | 5.81 |  | 0.031 |
|  |  | average tree girth | 3 | 63.1 | 5.92 |  | 0.029 |
