## Appendix D for "Habitat patch size and tree species richness shape the bird community in urban green spaces of rapidly urbanizing region of India"

Table D: Model description, scale, degrees of freedom (df) Akaike Information Criteria (AICc), relative support for hypothesis (AICc), of candidate models explaining find-foraging guild richness as a function of landscape and local variables during breeding season.

| Guild | Scale | Model | df | AICc | ΔAICc | weight |
| --- | --- | --- | --- | --- | --- | --- |
| Understory insectivore | *Landscape* | Area of urban green space | 2 | 76.1 | 0 | 0.888 |
|  |  | % barren+ % agriculture | 3 | 82.3 | 6.23 | 0.039 |
|  |  | Null model | 1 | 83.1 | 7.04 | 0.026 |
|  |  | % green | 2 | 83.5 | 7.43 | 0.022 |
|  |  | % builtup | 2 | 85.6 | 9.46 | 0.008 |
|  |  | % of open | 2 | 85.7 | 9.58 | 0.007 |
|  |  | % barren | 2 | 85.7 | 9.61 | 0.007 |
|  |  | % barren+% open | 3 | 88.7 | 12.55 | 0.002 |
|  | *Local* | tree richness | 2 | 74.9 | 0 | 0.707 |
|  |  | tree richness+shrub richness | 3 | 77.3 | 2.41 | 0.212 |
|  |  | shrub richness | 2 | 80.5 | 5.52 | 0.045 |
|  |  | Null model | 1 | 83.1 | 8.21 | 0.012 |
|  |  | average tree girth | 2 | 83.4 | 8.46 | 0.01 |
|  |  | shrub cover | 2 | 84.7 | 9.79 | 0.005 |
|  |  | tree crown cover | 2 | 85.3 | 10.41 | 0.004 |
|  |  | average tree height | 2 | 85.5 | 10.53 | 0.004 |
|  |  | tree crown cover+shrub cover | 3 | 87.3 | 12.39 | 0.001 |
| Sallying insectivore | *Landscape* | Area of urban green space | 2 | 60.7 | 0 | 0.42 |
|  |  | % barren | 2 | 62 | 1.25 | 0.225 |
|  |  | Null model | 1 | 63.4 | 2.68 | 0.11 |
|  |  | % green | 2 | 64.8 | 4.06 | 0.055 |
|  |  | % barren+% open | 3 | 64.8 | 4.09 | 0.054 |
|  |  | % barren+ % agriculture | 3 | 64.8 | 4.09 | 0.054 |
|  |  | % builtup | 2 | 64.9 | 4.18 | 0.052 |
|  |  | % of open | 2 | 66 | 5.26 | 0.03 |
|  | *Local* | tree richness | 2 | 60.4 | 0 | 0.37 |
|  |  | average tree girth | 2 | 61.5 | 1.16 | 0.207 |
|  |  | shrub cover | 2 | 63.3 | 2.9 | 0.087 |
|  |  | tree richness+shrub richness | 3 | 63.3 | 2.95 | 0.085 |
|  |  | Null Model | 1 | 63.4 | 3.02 | 0.082 |
|  |  | shrub richness | 2 | 64.2 | 3.81 | 0.055 |
|  |  | average tree height | 2 | 64.2 | 3.81 | 0.055 |
|  |  | tree crown cover | 2 | 65.3 | 4.94 | 0.031 |
|  |  | tree crown cover+shrub cover | 3 | 65.6 | 5.17 | 0.028 |
| Canopy insectivore | *Landscape* | Area of urban green space | 2 | 57.3 | 0 | 0.985 |
|  |  | Null model | 1 | 67.6 | 10.31 | 0.006 |
|  |  | % barren | 2 | 69.9 | 12.59 | 0.002 |
|  |  | % green | 2 | 69.9 | 12.64 | 0.002 |
|  |  | % barren+ % agriculture | 3 | 69.9 | 12.66 | 0.002 |
|  |  | % builtup | 2 | 70 | 12.76 | 0.002 |
|  |  | % of open | 2 | 70 | 12.77 | 0.002 |
|  |  | % barren+% open | 3 | 72.6 | 15.34 | 0 |
|  | *Local* | tree richness | 2 | 62.1 | 0 | 0.32 |
|  |  | shrub cover | 2 | 62.2 | 0.03 | 0.316 |
|  |  | tree richness+shrub richness | 3 | 63.8 | 1.65 | 0.14 |
|  |  | shrub richness | 2 | 64.3 | 2.19 | 0.107 |
|  |  | tree crown cover+shrub cover | 3 | 65.1 | 2.99 | 0.072 |
|  |  | Null Model | 1 | 67.6 | 5.46 | 0.021 |
|  |  | average tree girth | 2 | 68.7 | 6.58 | 0.012 |
|  |  | average tree height | 2 | 70.1 | 7.96 | 0.006 |
|  |  | tree crown cover | 2 | 70.1 | 8.01 | 0.006 |
| Granivore | *Landscape* | % barren | 2 | 60.1 | 0 | 0.337 |
|  |  | % barren+ % agriculture | 3 | 61.3 | 1.19 | 0.186 |
|  |  | Area of urban green space | 2 | 62.1 | 2 | 0.124 |
|  |  | Null model | 1 | 62.4 | 2.26 | 0.109 |
|  |  | % barren+% open | 3 | 62.7 | 2.56 | 0.094 |
|  |  | % builtup | 2 | 63.3 | 3.15 | 0.07 |
|  |  | % green | 2 | 64 | 3.8 | 0.05 |
|  |  | % of open | 2 | 65 | 4.85 | 0.03 |
|  | *Local* | average tree height | 2 | 58 | 0 | 0.586 |
|  |  | tree richness | 2 | 61.5 | 3.5 | 0.102 |
|  |  | average tree girth | 2 | 61.6 | 3.63 | 0.095 |
|  |  | shrub richness | 2 | 62.4 | 4.4 | 0.065 |
|  |  | Null Model | 1 | 62.4 | 4.4 | 0.065 |
|  |  | tree richness+shrub richness | 3 | 63.9 | 5.88 | 0.031 |
|  |  | tree crown cover | 2 | 64 | 5.98 | 0.029 |
|  |  | shrub cover | 2 | 64.8 | 6.8 | 0.02 |
|  |  | tree crown cover+shrub cover | 3 | 66.8 | 8.76 | 0.007 |
| Nectar insectivore | *Landscape* | Null model | 1 | 60.9 | 0 | 0.33 |
|  |  | Area of urban green space | 2 | 62 | 1.05 | 0.195 |
|  |  | % green | 2 | 62.9 | 1.96 | 0.123 |
|  |  | % builtup | 2 | 62.9 | 1.99 | 0.122 |
|  |  | % of open | 2 | 63.5 | 2.53 | 0.093 |
|  |  | % barren | 2 | 63.5 | 2.54 | 0.092 |
|  |  | % barren+ % agriculture | 3 | 66.3 | 5.35 | 0.023 |
|  |  | % barren+% open | 3 | 66.4 | 5.45 | 0.022 |
|  | *Local* | Null Model | 1 | 60.9 | 0 | 0.254 |
|  |  | shrub cover | 2 | 61.8 | 0.83 | 0.168 |
|  |  | average tree height | 2 | 62.4 | 1.4 | 0.126 |
|  |  | average tree girth | 2 | 62.6 | 1.61 | 0.114 |
|  |  | tree richness | 2 | 62.8 | 1.85 | 0.101 |
|  |  | tree crown cover | 2 | 63 | 2.01 | 0.093 |
|  |  | shrub richness | 2 | 63.5 | 2.56 | 0.071 |
|  |  | tree crown cover+shrub cover | 3 | 64.2 | 3.22 | 0.051 |
|  |  | tree richness+shrub richness | 3 | 65.7 | 4.75 | 0.024 |
| Fruit-seed-nectar-insectivore | *Landscape* | Area of urban green space | 2 | 68.1 | 0 | 0.301 |
|  |  | Null model | 1 | 68.2 | 0.08 | 0.289 |
|  |  | % of open | 2 | 70.6 | 2.5 | 0.086 |
|  |  | % builtup | 2 | 70.8 | 2.65 | 0.08 |
|  |  | % green | 2 | 70.8 | 2.67 | 0.079 |
|  |  | % barren | 2 | 70.8 | 2.67 | 0.079 |
|  |  | % barren+ % agriculture | 3 | 71.2 | 3.06 | 0.065 |
|  |  | % barren+% open | 3 | 73.6 | 5.49 | 0.019 |
|  | *Local* | tree richness | 2 | 67.2 | 0 | 0.336 |
|  |  | Null Model | 1 | 68.2 | 1.02 | 0.202 |
|  |  | shrub cover | 2 | 69.5 | 2.3 | 0.106 |
|  |  | tree richness+shrub richness | 3 | 70.2 | 2.96 | 0.077 |
|  |  | shrub richness | 2 | 70.2 | 2.98 | 0.076 |
|  |  | average tree girth | 2 | 70.4 | 3.2 | 0.068 |
|  |  | tree crown cover | 2 | 70.8 | 3.59 | 0.056 |
|  |  | average tree height | 2 | 70.8 | 3.61 | 0.055 |
|  |  | tree crown cover+shrub cover | 3 | 72.5 | 5.27 | 0.024 |
| Fruit-seed-nectar | *Landscape* | Null model | 1 | 49 | 0 | 0.281 |
|  |  | Area of urban green space | 2 | 49.5 | 0.42 | 0.227 |
|  |  | % barren | 2 | 50.6 | 1.54 | 0.13 |
|  |  | % builtup | 2 | 51.2 | 2.13 | 0.097 |
|  |  | % green | 2 | 51.6 | 2.52 | 0.08 |
|  |  | % barren+ % agriculture | 3 | 51.6 | 2.53 | 0.079 |
|  |  | % of open | 2 | 51.6 | 2.59 | 0.077 |
|  |  | % barren+% open | 3 | 53.6 | 4.51 | 0.029 |
|  | *Local* | Null Model | 1 | 49 | 0 | 0.289 |
|  |  | tree richness | 2 | 49.9 | 0.82 | 0.192 |
|  |  | shrub richness | 2 | 51.1 | 2.09 | 0.102 |
|  |  | shrub cover | 2 | 51.3 | 2.3 | 0.092 |
|  |  | tree crown cover | 2 | 51.4 | 2.35 | 0.089 |
|  |  | average tree height | 2 | 51.4 | 2.4 | 0.087 |
|  |  | average tree girth | 2 | 51.5 | 2.5 | 0.083 |
|  |  | tree richness+shrub richness | 3 | 52.8 | 3.8 | 0.043 |
|  |  | tree crown cover+shrub cover | 3 | 54.1 | 5.04 | 0.023 |
| Raptor | *Landscape* | Area of urban green space | 2 | 47.4 | 0 | 0.342 |
|  |  | Null model | 1 | 48.2 | 0.78 | 0.231 |
|  |  | % green | 2 | 49.8 | 2.39 | 0.103 |
|  |  | % barren | 2 | 50.3 | 2.88 | 0.081 |
|  |  | % barren+ % agriculture | 3 | 50.3 | 2.89 | 0.08 |
|  |  | % of open | 2 | 50.5 | 3.04 | 0.075 |
|  |  | % builtup | 2 | 50.7 | 3.25 | 0.067 |
|  |  | % barren+% open | 3 | 53.1 | 5.65 | 0.02 |
|  | *Local* | Null Model | 1 | 48.2 | 0 | 0.252 |
|  |  | tree richness | 2 | 48.8 | 0.6 | 0.187 |
|  |  | average tree height | 2 | 49.4 | 1.19 | 0.139 |
|  |  | shrub cover | 2 | 49.8 | 1.63 | 0.112 |
|  |  | shrub richness | 2 | 50.4 | 2.18 | 0.085 |
|  |  | tree crown cover | 2 | 50.4 | 2.2 | 0.084 |
|  |  | average tree girth | 2 | 50.8 | 2.59 | 0.069 |
|  |  | tree richness+shrub richness | 3 | 51.8 | 3.58 | 0.042 |
|  |  | tree crown cover+shrub cover | 3 | 52.4 | 4.19 | 0.031 |
| Omnivore | *Landscape* | Area of urban green space | 2 | 62.5 | 0 | 0.753 |
|  |  | Null model | 1 | 66.8 | 4.29 | 0.088 |
|  |  | % of open | 2 | 68.1 | 5.64 | 0.045 |
|  |  | % barren | 2 | 69.1 | 6.56 | 0.028 |
|  |  | % green | 2 | 69.1 | 6.6 | 0.028 |
|  |  | % builtup | 2 | 69.3 | 6.8 | 0.025 |
|  |  | % barren+ % agriculture | 3 | 69.5 | 7.04 | 0.022 |
|  |  | % barren+% open | 3 | 71 | 8.49 | 0.011 |
|  | *Local* | tree richness | 2 | 65.1 | 0 | 0.332 |
|  |  | Null Model | 1 | 66.8 | 1.7 | 0.142 |
|  |  | shrub richness | 2 | 66.8 | 1.76 | 0.138 |
|  |  | shrub cover | 2 | 67 | 1.89 | 0.129 |
|  |  | tree richness+shrub richness | 3 | 67.7 | 2.58 | 0.091 |
|  |  | average tree girth | 2 | 68.8 | 3.73 | 0.051 |
|  |  | average tree height | 2 | 69.1 | 4.03 | 0.044 |
|  |  | tree crown cover | 2 | 69.2 | 4.16 | 0.042 |
|  |  | tree crown cover+shrub cover | 3 | 69.8 | 4.74 | 0.031 |
| Trunk-bark forager | *Landscape* | Null model | 1 | 37.4 | 0 | 0.292 |
|  |  | Area of urban green space) | 2 | 38.3 | 0.83 | 0.193 |
|  |  | % barren | 2 | 38.9 | 1.49 | 0.138 |
|  |  | % builtup | 2 | 39.4 | 1.94 | 0.11 |
|  |  | % of open | 2 | 39.5 | 2.08 | 0.103 |
|  |  | % green | 2 | 40 | 2.54 | 0.082 |
|  |  | % barren+% open | 3 | 41.2 | 3.77 | 0.044 |
|  |  | % barren+ % agriculture | 3 | 41.5 | 4.08 | 0.038 |
|  | *Local* | tree richness | 2 | 35.9 | 0 | 0.41 |
|  |  | Null Model | 1 | 37.4 | 1.51 | 0.193 |
|  |  | tree richness+shrub richness | 3 | 38.8 | 2.88 | 0.097 |
|  |  | shrub richness | 2 | 39.6 | 3.66 | 0.066 |
|  |  | average tree girth | 2 | 39.8 | 3.86 | 0.06 |
|  |  | shrub cover | 2 | 39.9 | 3.99 | 0.056 |
|  |  | tree crown cover | 2 | 40 | 4.05 | 0.054 |
|  |  | average tree height | 2 | 40 | 4.1 | 0.053 |
|  |  | tree crown cover+shrub cover | 3 | 42.8 | 6.93 | 0.013 |
| Frugivore insectivore | *Landscape* | % barren | 2 | 64.2 | 0 | 0.266 |
|  |  | Area of urban green space | 2 | 64.4 | 0.2 | 0.24 |
|  |  | % barren+ % agriculture | 3 | 65.7 | 1.51 | 0.125 |
|  |  | Null model | 1 | 65.8 | 1.55 | 0.123 |
|  |  | % builtup | 2 | 65.9 | 1.73 | 0.112 |
|  |  | % barren+% open | 3 | 67.1 | 2.92 | 0.062 |
|  |  | % of open | 2 | 68.1 | 3.87 | 0.039 |
|  |  | % green | 2 | 68.3 | 4.13 | 0.034 |
|  | *Local* | tree richness | 2 | 65.2 | 0 | 0.296 |
|  |  | Null Model | 1 | 65.8 | 0.6 | 0.219 |
|  |  | average tree height | 2 | 67.4 | 2.22 | 0.098 |
|  |  | shrub richness | 2 | 67.5 | 2.35 | 0.091 |
|  |  | tree crown cover | 2 | 67.8 | 2.62 | 0.08 |
|  |  | average tree girth | 2 | 68.1 | 2.96 | 0.067 |
|  |  | tree richness+shrub richness | 3 | 68.1 | 2.99 | 0.066 |
|  |  | shrub cover | 2 | 68.2 | 3.08 | 0.064 |
|  |  | tree crown cover+shrub cover | 3 | 70.7 | 5.5 | 0.019 |
| Ground insectivore | *Landscape* | % barren | 2 | 58.4 | 0 | 0.307 |
|  |  | Area of urban green space) | 2 | 59.3 | 0.81 | 0.205 |
|  |  | Null model | 1 | 60 | 1.56 | 0.14 |
|  |  | % green | 2 | 60.8 | 2.41 | 0.092 |
|  |  | % builtup | 2 | 61.3 | 2.88 | 0.073 |
|  |  | % barren+ % agriculture | 3 | 61.3 | 2.9 | 0.072 |
|  |  | % barren+% open | 3 | 61.4 | 2.92 | 0.071 |
|  |  | % of open | 2 | 62.5 | 4.09 | 0.04 |
|  | *Local* | average tree height | 2 | 55.4 | 0 | 0.486 |
|  |  | shrub richness | 2 | 57.7 | 2.24 | 0.158 |
|  |  | average tree girth | 2 | 57.9 | 2.41 | 0.146 |
|  |  | tree richness | 2 | 59.4 | 3.98 | 0.067 |
|  |  | Null Model | 1 | 60 | 4.56 | 0.05 |
|  |  | tree richness+shrub richness | 3 | 60.1 | 4.67 | 0.047 |
|  |  | tree crown cover | 2 | 61.3 | 5.86 | 0.026 |
|  |  | shrub cover | 2 | 62.6 | 7.13 | 0.014 |
|  |  | tree crown cover+shrub cover | 3 | 64.3 | 8.82 | 0.006 |
